# Do allochthonous flows explain deviations from the Redfield ratio in lakes?

**DOI:** 10.1101/2025.07.06.663341

**Authors:** Benoît Pichon, Tanguy Daufresne, Frédéric Guichard, Isabelle Gounand

**Author notes:** Contributed equally (Tanguy Daufresne, Isabelle Gounand).

## Abstract

Lakes or streams show strong deviations of their stoichiometry compared to the Redfield ratio measured in oceans. Allochthonous inflows of resources might contribute to those deviations directly, by changing composition of detritus in ecosystems, but also indirectly, by shaping local community dynamics and stoichiometric constrains within aquatic ecosystems. Here, we developed a stoichiometric model to understand those direct and indirect mechanisms through which allochthonous inflows affect seston stoichiometry. Our results emphasize that increasing allochthonous inflows promotes heterotrophic functioning, and relax decomposer’ carbon limitation. This release of stoichiometric constrain of decomposers *(i)* destabilizes the aquatic ecosystem by promoting competition between phytoplankton and decomposers, *(ii)* decreases the ability of the lake to regulate allochthonous flows, and *(iii)* push seston stoichiometry away from the Redfield ratio. Our study emphasize how the quantity and the stoichiometry of inflows shape community dynamics and the elemental constraints within the ecosystem, and open perspectives for stoichiometry at the landscape extent.

## Introduction

Perhaps, one of the most intriguing patterns in biogeochemistry is the relative stability of carbon (C), nitrogen (N) and phosphorus (P) ratios in aquatic ecosystems (hereafter called stoichiometric homeostasis). Albert Redfield was among the first to observe an atomic ratio of C:N:P in oceans around 106:16:1, which he related to the average stoichiometric ratios of phytoplankton (Redfield, 1934, 1958). The detail of this regulation was later shown to be based on the competition between planktonic species (Tyrrell, 1999; Sterner & Elser, 2002). Specifically, when nitrogen supply is limited, the C:N:P ratio in oceans is maintained by nitrogen fixers adding nitrogen by N_2_ fixation, while when phosphorus is limiting, N-fixers become less competitive, and non-fixers species become dominant and deplete nitrogen. Later, deviations of seston (*i.e.,* plankton and particulate detritus) stoichiometry from the Redfield ratio in oceans have been explained by planktonic species composition, species-specific stoichiometric constraints (quotas), nitrogen or phosphorus of plankton, or differences in nutrient allocations among species (Klausmeier et al., 2004; Weber & Deutsch, 2010; Martiny et al., 2013). Yet, these explanations do not transfer well to lakes where large deviations from the Redfield ratio are observed (Sterner et al., 2008). To explain lake stoichiometry, early studies have therefore proposed that within-lake processes such as denitrification or respiration (Hecky et al., 1993), or lake size characteristics (Sterner et al., 2008) could contribute to the observed deviations. However, external influences, particularly allochthonous inflows, may also play a significant role. Although ecosystems were once viewed as relatively closed units (O’Neill, 2001), they are open to exchanges of resources, organisms, energy, and information (Polis et al., 1997; Massol et al., 2011; Bartels et al., 2012; Gounand et al., 2018). Examples include terrestrial ecosystems exporting carbon to aquatic ecosystems (*e.g.,* from leaves windblown and sinking; Pace et al., 2004), or the sinking of nutrients and feces into benthic areas in lakes (Bergströ m et al., 2003). Here, we collected from the literature stoichiometric ratios of allochthonous inflows exported from forests and grasslands to lakes and streams (Appendix S1 for details), and we show that elemental ratios of resources exported by terrestrial ecosystems vary a lot, from 3.8-169 for C:N, 2-7198 for C:P, and 3-100 for N:P depending on the type of inflow (Table S4), consistent with findings by (Bartels et al., 2012; Sitters et al., 2015). The stoichiometric ratios of many subsidy types deviate from the Redfield ratio (Fig. 1). Notably, terrestrial inflows from plant litter or leaves had on average higher C:P and C:N ratios than the Redfield ratio. Carbon rich forest inflows might therefore push aquatic ecosystems more quickly to nutrient limitation compared to more nutrient-rich subsidies (*e.g.,* carcasses, fertilizer leaching).

**Figure 1:**
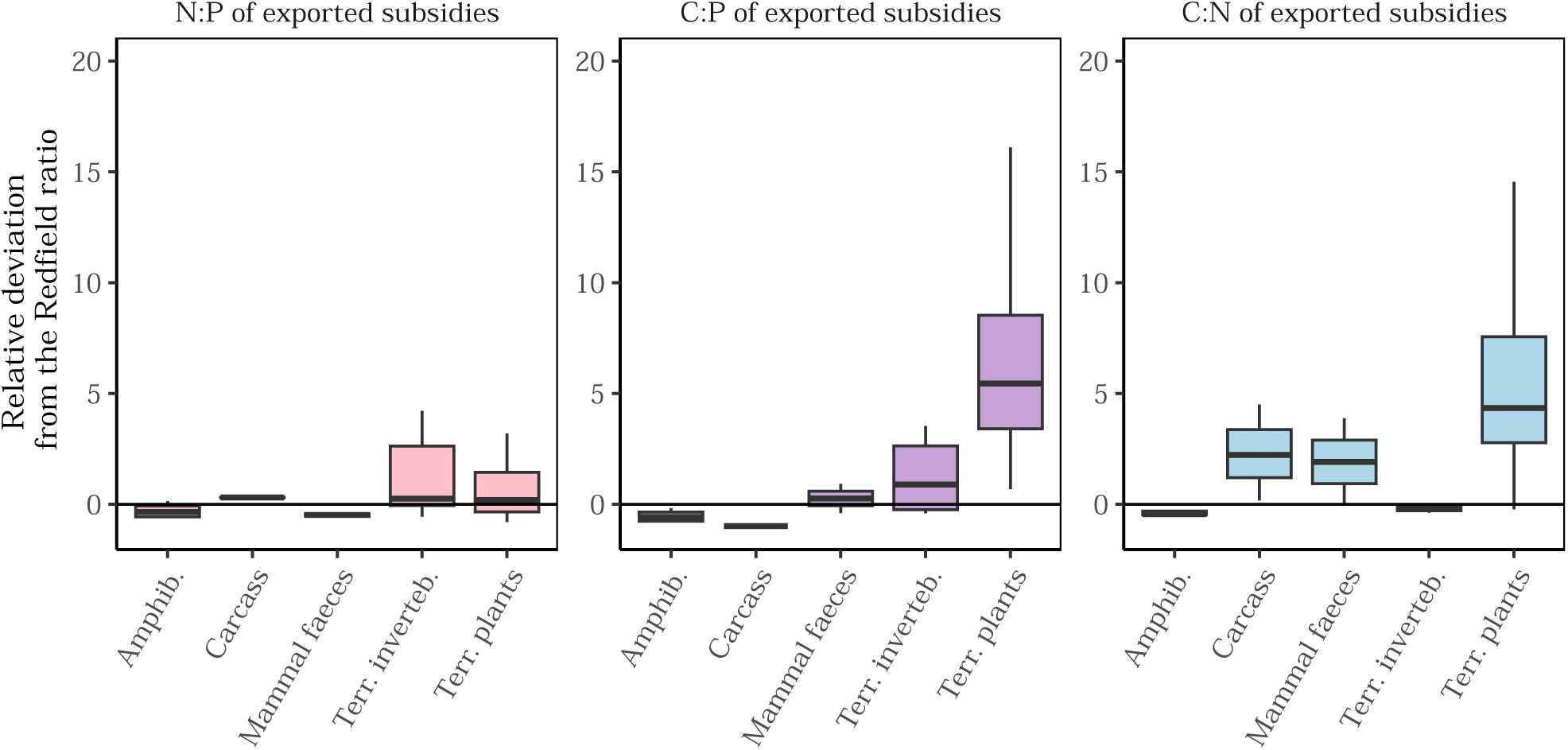
Deviations from the Redfield ratio of stoichiometry of terrestrial allochthonous inflows exported in lakes and streams. Relative deviations of elemental ratios (C:P, C:N, and N:P) of the resource flows to the Redfield ratio (*X_obs_*− *X_Red f ield_*)/(*X_Red f ield_*) for different type of materials exported to lakes and streams: Fresh., freshwater; Terr., terrestrial; amphib., amphibians and inverteb., invertebrates. Boxplots present the median surrounded by the first and third quartiles, and whiskers are not shown. We refer to Appendix S1 for details on the paper used to produce these figures and on the general methodology.

Allochthonous inflows might affect lake stoichiometry both by directly altering the elemental composition of the seston, and indirectly by inducing changes in community structure, inorganic nutrient pools, and organismal stoichiometric demands. Large amounts of allochthonous flows can leave a strong imprint on lake seston stoichiometry, especially in small lakes which have lower dilution capacity then larger lakes. This could explain why lakes with larger areas show lower deviation from the Redfield ratio compared to smaller ones (Hecky et al., 1993; Sterner et al., 2008). Beyond mass effects, later studies emphasized the importance of land-use and land-cover of the regional landscape (*e.g.,* agricultural *versus* forested areas) on lake stoichiometry (Hessen et al., 2009; Vanni et al., 2011; Prater et al., 2017; Soranno et al., 2019). Land-use from neighboring ecosystems can export nutrients and detritus of different stoichiometries, which directly changes the elemental ratios in the seston they integrate (Sterner et al., 2008). For example, allochthonous inflows from pasture areas have lower N:P compared to those from agricultural crops (Arbuckle & Downing, 2001), while allochthonous inflows from forested areas tend to be richer in carbon as compared to those from agricultural areas (Pichon et al., 2023). As such, neighboring crops or forests were identified as among the main drivers of N:P ratios in US lakes Collins et al. (2017). In addition to these direct pathways, the indirect effects of allochthonous inflows, mediated through changes in nutrient stocks, biotic interactions, and stoichiometric constraints, further shape seston composition (Pichon et al., 2023; Osakpolor et al., 2023). Hitherto, despite growing hypotheses to explain deviations from the Redfield ratio, a mechanistic understanding of how landscape context drives direct and indirect effects on seston C:N:P ratios remains limited.

Understanding the elemental ratios of lakes’ seston requires unveiling the interactions of decomposers and phytoplankton and their consequences on the stoichiometric homeostasis of the seston, which has been overlooked so far (Tyrrell, 1999; Hessen et al., 2003). Decomposers participate in nutrient cycling through decomposition and mineralization processes (McClaugherty et al., 1984), and are therefore key cogs of the recycling machinery. In response to allochthonous inflows of nutrients, decomposition can be stimulated, which fosters carbon losses from aquatic ecosystems (Rosemond et al., 2015). In addition, the contribution of lake decomposers to observed sestonic elemental ratios in lakes might be important since they have, on average, lower N:C and P:C contents compared to phytoplankton (Buchkowski et al., 2019; Sterner & Elser, 2002), and can be particularly abundant in some lakes with high riparian cover or receiving large amounts of allochthonous carbon (Bergströ m et al., 2003). In fact, a recent study found that clear lakes had lower N:P and C:P ratios while having higher C:N ratios compared to turbid ones (Ginger et al., 2017). For instance, Bergströ m et al., 2003 found that C addition in lakes led to higher abundance of bacteria and to lower abundance of autotrophs, while the inverse was found with N or P addition. Consequently, we hypothesize that allochthonous inflows of resources may modulate the level of heterotrophy, *i.e.*, the ratio of carbon respired to carbon fixed, and leave an imprint on the sestonic C:N:P ratios of lakes (see Fig. 1a). This might be especially true in small lakes, where stronger deviations from the Redfield ratio are observed compared to large lakes (Sterner et al., 2008).

Decomposers not only are at the basis of the recycling loop in ecosystems (Veldhuis et al., 2018), they are also involved in complex sets of interactions with the phytoplankton: decomposers and phytoplankton compete for nutrients, but are also involved in an indirect mutualism through the mineralization of nutrients by the decomposers and the production of detritus by the phytoplankton therefore producing resources for their respective competitor (Daufresne & Loreau, 2001; Danger et al., 2007b). In lakes or streams, the coexistence of decomposers and phytoplankton is maintained because bacteria is generally limited (or co-limited) by carbon and another nutrient (Bergströ m et al., 2003; Daufresne et al., 2008; but see Hessen et al., 1994), and by stratification, which induces a spatial segregation. The functional implication of these interactions have been investigated in small experimental settings (Danger et al., 2007b), and in models to understand ecosystem (Cherif & Loreau, 2013; Buchkowski et al., 2019) and landscape functioning (Pichon et al., 2023), but has not been yet investigated in a spatial context with allochthonous inflows from neighboring ecosystems.

We therefore ask the following question: how do allochthonous inflows of resources modulate the elemental ratios of the seston in lake ecosystems through changes in community composition of decomposers and phytoplanktonTo study this question, we developed a stoichiometric ecosystem model for lakes with high riparian cover and open to allochthonous inflows of resources (Fig. 1). The model represents the interactions between phytoplankton–either nitrogen fixers or non-fixers– and decomposers (heterotrophic bacteria). The model is analyzed in the context of lakes but is general enough to model the stoichiometric interactions between phytoplankton and decomposers in oceans or streams as well. We analyzed the model to understand how allochthonous inflows modulate local community dynamics, lake heterotrophy, as well as the scaling of C:N:P ratios and their deviations from the Redfield ratio. Our results reveal that both quality and quantity of allochthonous inflows interact with local stoichiometric constraints of phytoplankton and decomposers to drive the level of heterotrophy and the stoichiometric ratios of seston in lakes.

## Material and methods

### Model description

#### A stoichiometric model of aquatic ecosystems

We built a stoichiometric model of aquatic ecosystems that couples together the dynamics of carbon, nitrogen, and phosphorus (Fig. 2). The lake ecosystem is composed of three types of microbial functional groups: autotrophic phytoplankton with nitrogen fixers (*F*) and non-fixers (*O*), and heterotrophic decomposers (*B*), together representing basal biomass production in the ecosystem. Herbivory is implicitly represented in phytoplankton mortality (see below). The ecosystem has a detritus (*D*) pool and two inorganic nutrient ones: nitrogen and phosphorus (*N*, *P* respectively). The detritus pool together with the three functional groups represent the seston in our model (white rectangle in Fig. 2). We followed the carbon content of the detritus pool and three functional groups (*i.e.*, *D_C_*, *F_C_*, *O_C_*, and *B_C_*). We derived the nitrogen and phosphorus content of functional groups (*i.e.*, *F_N_*, *F_P_*, *O_N_*, *O_P_*, which equations are omitted here, and *B_N_*, *B_P_*) by assuming a stoichiometric homeostasis with fixed molar nitrogen (*resp.* phosphorus) to carbon ratio N:C *α_X_* (*resp.* P:C, *β_X_*) where *X* ∈ {*F*, *O*, *B*}. We acknowledge this simplifying assumption for both phytoplankton types, as they can partly adjust their stoichiometry depending on nutrient availability (Sterner & Elser, 2002), and decomposers are not always homeostatic (Scott et al., 2012). Detritus pool has its stoichiometric ratios *α_D_* and *β_D_* that dynamically change depending on the detritus produced by the different organisms and on the allochthonous inflows (see below).

**Figure 2:**
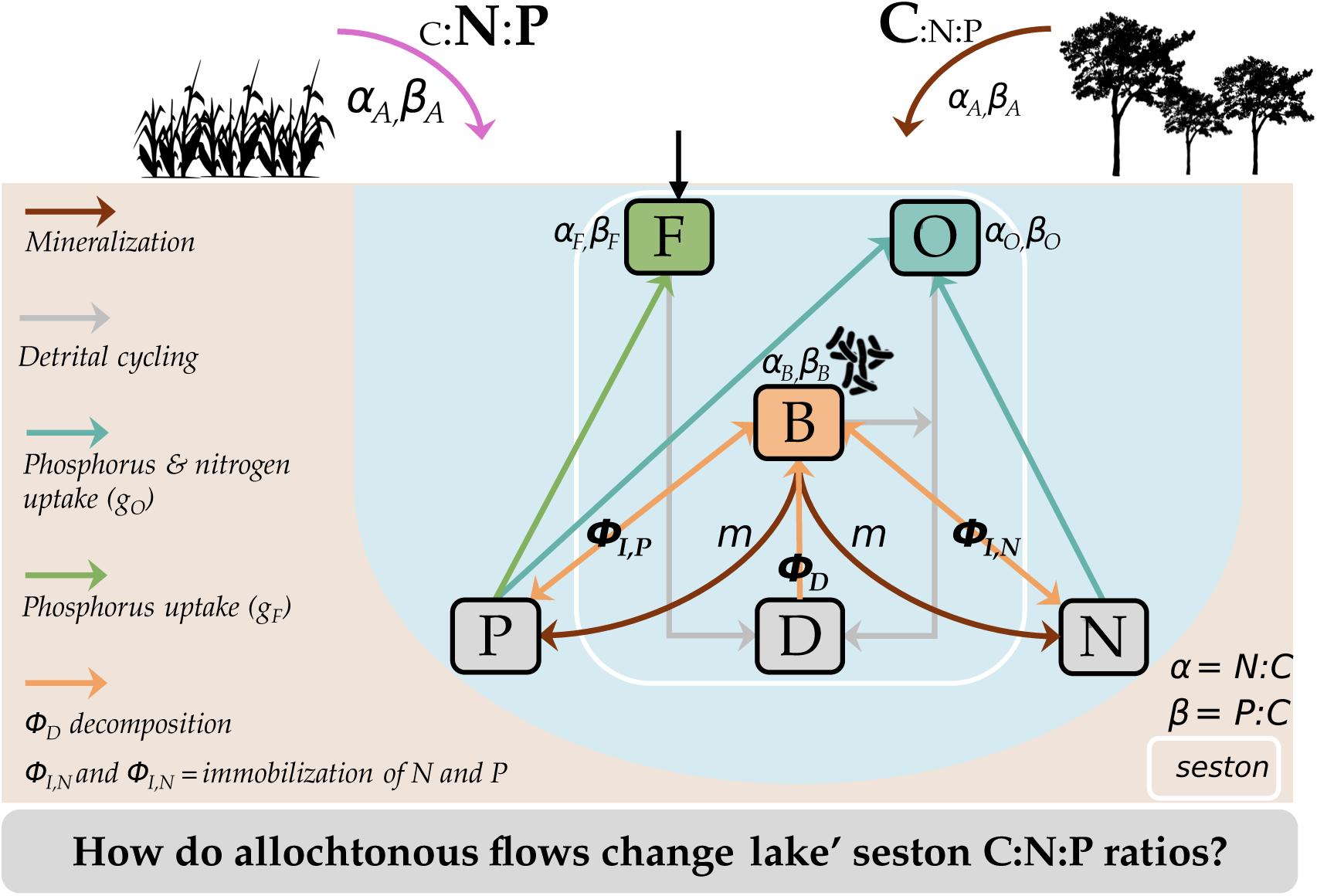
A carbon, nitrogen, and phosphorus model of aquatic ecosystems open to allochthonous inflows. To explore these hypotheses and elucidate some mechanisms, we built a stoichiometric model of lake dynamics open to allochthonous inflows. The aquatic ecosystem is composed of inorganic nutrients (phosphorus *P* and nitrogen *N*), detritus (*D*), and three functional groups of organisms: phytoplankton that can either fix nitrogen (fixers, *F*) or not (non-fixers, *O*), and decomposers (*B*). Each of the three functional groups maintains fixed stoichiometric ratios N:C (*α*) and P:C (*β*). Fixers fix nitrogen via biological fixation and are therefore exclusively phosphorus-limited. Non-fixers by contrast can be either nitrogen and phosphorus-limited depending on the relative availability of nutrients. Decomposers have three possible limitations: nitrogen, phosphorus, or carbon. Their limitation sets which flow (decomposition of detritus (*ϕ_D_*) or immobilization of nitrogen or phosphorus (*ϕ_I_*_,*N*_, *ϕ_I_*_,*P*_)) constrains the others so that decomposers maintain a constant stoichiometry (Appendix A). The other symbols are used in the model equations: mineralization rate of decomposers (*m*), fixers growth function (*g_F_*), non-fixers growth function (*g_O_*). The aquatic ecosystem is open to allochthonous inflows with specific stoichiometric ratios *α_A_* and *β_A_*.

#### Decomposers and phytoplankton dynamics

Heterotrophic decomposers primarily feed on detritus (Fig. 2), and can be limited by carbon, nitrogen, or phosphorus. Since the elemental ratios of the detritus pool do not always match their requirements (*e.g., β_B_* ≠ *β_D_* for P:C ratio) due to the different stoichiometry of the detritus produced by phytoplankton or due to allochthonous inflows, decomposers take-up (immobilization) or excrete nitrogen and phosphorus (mineralization) to maintain their homeostasis (Daufresne & Loreau, 2001). In this model, the limitations of decomposers and non-fixers are emergent characteristics of the detritus dynamics in the system (see Appendix A for the formula to determine decomposer’ limitation). When decomposers are *N*− (*resp. P*−) limited, the decomposition flux (*i.e.,* consumption of detritus, *ϕ_D_*) is constrained by nitrogen (*resp.* phosphorus) immobilization’ (*i.e.,* nutrient uptake *ϕ_I_*_,*N*_ or *ϕ_I_*_,*P*_) whereas, when decomposers are *C*−limited, decomposition determines nutrient uptake.

By contrast, nitrogen fixers are exclusively limited by phosphorus for growth since they can fix nitrogen and carbon, while non-fixers can be limited by nitrogen or phosphorus depending on their relative availability (modelled with a Liebig law of the minimum). Both phytoplankton types maintain their stoichiometric homeostasis by adjusting their nutrient intake, which are given by type-I functional responses *g_F_*(*P*) = *µ_F_P* and *g_O_*(*N*, *P*) = *µ_O_*min(*N*, *^βO^ P*) for fixers and non-fixers respectively.

Since our primary goal is to understand how allochthonous inflows modulate local interactions between the three functional groups and ultimately the sestonic elemental ratios, we assumed a self-regulation term for each functional group (*s_X_X*^2^). This term allows to *(i)* maintain coexistence despite potential similar limitations (Simha et al., 2022), *(ii)* stabilize the dynamics (Barbier & Loreau, 2019), and *(iii)* implicitly account for top-down control by zooplankton in the aquatic ecosystem. In a lake for instance, decomposers and phytoplankton can coexist despite varying levels of bacteria production and allochthonous inflows since water depth or water column stratification might limit cases of strict competitive exclusion (Bergströ m et al., 2003).

Recycling occurs through the production of detritus at a rate *d_X_*by each functional group *X*, fuelling the detritus pool, as well as by the mineralization of detritus into nitrogen and phosphorus at a rate *m* by decomposers.

#### Allochthonous flows of nutrients, carbon and phosphorus

The ecosystem is open to autochthonous fluxes, *I_N_*, *I_P_*, *I_D_*, fueling nitrogen, phosphorus and detritus stocks respectively. There are also losses at rates *l_N_*, *l_P_*, *l_D_*, by denitrification or sinking, not explicitly accounted for in this model. Allochthonous detritus do not always have C:N:P ratios matching the ones of local detritus, and therefore we define specific N:C and P:C ratios for allochthonous detritus (*α_A_*, *β_A_* respectively). Here are the ecological dynamics given that fixers and non-fixers are homeostatic (*F_N_* = *α_F_F_C_*):

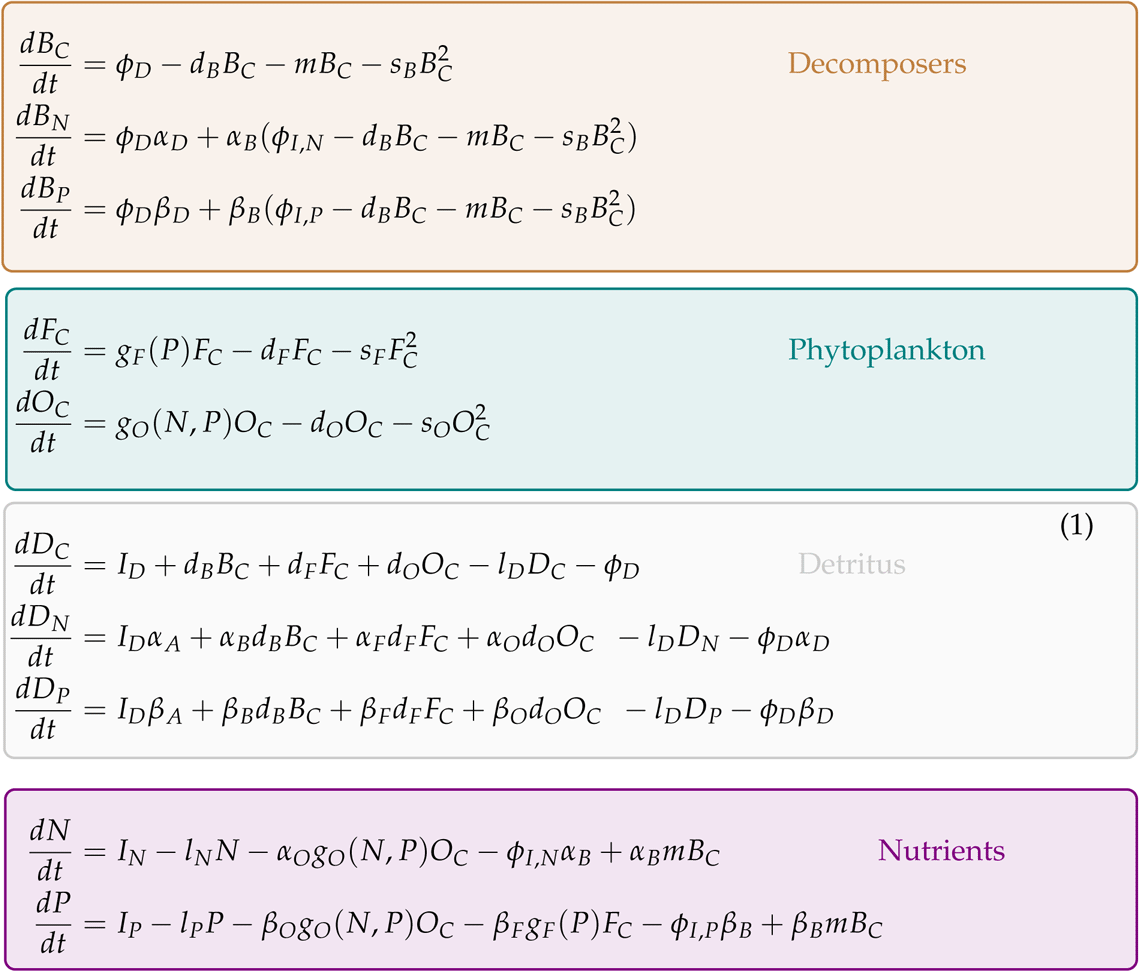

### Quantifying indirect effects, seston stoichiometric ratio and level of heterotrophy

Since we want to test how the interaction between allochthonous inflows, and interactions between decomposers and phytoplankton mediate elemental ratios in the aquatic ecosystem, we quantified *(i)* the indirect interactions between the different functional groups, *(ii)* the stoichiometric ratios in the seston and their partitioning between detritus pool and functional groups (see below).

#### Quantifying indirect interactions

We measured the effect of functional group *j* (*j* ∈ {*F*, *O*, *B*}) on *i* as their indirect interactions *I_ij_*. Indirect effects change depending on the limitation of the different functional groups, their production of detritus and the level of mineralization of detritus by decomposers. Following Nakajima & Higashi (1995), we computed the net effect (direct + indirect) as the effect of the increase of one density unit in functional group *j* on functional group *i*’s growth rate (”abundance to inflow” following Nakajima & Higashi, 1995 denomination). Since there are no direct interactions between functional groups, the indirect effects were derived from the inverse of the Jacobian matrix (sensitivity matrix *S* = *J*^−1^) at equilibrium. The indirect effect of functional group *j* on *i* integrates effects across all paths of the system, except the ones that loop on functional group *i* or *j*, and is given by the following formula:

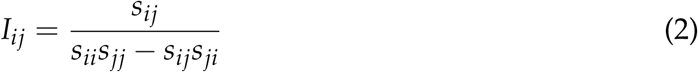

We note that using only the indirect effect without controlling for paths looping on the functional group *i* and *j* (*i.e., I_ij_* = *s_ij_*) gives qualitatively similar information about indirect competition or facilitation between functional groups (Fig. S1). We also computed a complementary metric, rooted in facilitation studies (Díaz-Sierra et al., 2017), to quantify how the density of each functional group changed with the presence of others, which we detail in Appendix A. This allows identifying scenarios where a given functional group may not persist in the presence of the others (*i.e.,* facilitation; or inversely competition).

#### Aquatic ecosystem production and C:N:P in the seston

We measured the N:C ratio of the seston by taking the ratio of the total nitrogen content in the detritus, phytoplankton and decomposers over their total carbon content (and similarly for the P:C ratio). We also defined the level of heterotrophy of the lake as the fraction of the density of decomposers (in carbon unit) divided by the cumulated density of phytoplankton and decomposers.

### Scenarios of inflows

We focus on the ecosystem dynamics under sustained increase in allochthonous inflows of detritus, *I_D_*, with different N:C and P:C ratios (*α_A_*, *β_A_*respectively) to account for both the quantity and quality of allochthonous inflows. While the quantity allows mimicking different lake sizes, with smaller lakes receiving proportionally more inflows than larger ones, the stoichiometry of allochthonous inflows allows testing different types of land-use from neighboring ecosystems. Specifically, we considered four scenarios of quality for the allochthonous inflows: nutrient-rich flows (*e.g.,* from agricultural areas with high P:C and high N:C), nitrogen-rich flows (*e.g.,* from agricultural crops with low P:C but high N:C), phosphorus rich flows (*e.g.,* from pasture with high P:C but low N:C), and carbon-rich flows (*e.g.,* from forested areas with plant litter with low P:C and low N:C, Box 1). These four scenarios allow investigating the different cases of decrease or increase of detritus N:C and P:C ratios in lakes by allochthonous inflows (see Fig. S2). We chose parameters that allow coexistence as a baseline and used the literature to inform realistic stoichiometric ratios of the different functional groups (Appendix). Decomposers have higher N:C and P:C than phytoplankton in our analysis following (Cleveland & Liptzin, 2007; Buchkowski et al., 2019; *α_B_ > α_F_*_,*O*_, *β_B_ > β_F_*_,*O*_). As such, detritus were poorer in nutrients as compared to the elemental constraints of decomposers (*α_D_ < α_B_*, *β_D_ < β_B_*). For simplicity, we did not implement differences in ratios among phytoplankton groups (*α_F_* = *α_O_*, *β_F_* = *β_O_*), but rather, we assumed that non-fixer phytoplankton were more competitive than fixers for phosphorus (*µ_O_> µ_F_*). Non-fixers remained nitrogen-limited on the parameters tested. All parameters used as well as their ranges of variation are made available in Appendix A. We performed simulations using *DifferentialEquation* package in Julia (version 1.7.3), and analysed them in R (version 4.4.1).

## Results

In the main text we present the results with the three functional groups but we refer to Appendix C to qualitatively similar results when only considering non-fixers and decomposers, or fixers and decomposers.

### Moderate allochthonous detritus promotes heterotrophy

Before inducing a change in limitation, increasing allochthonous detritus promotes heterotrophic production relative to primary production (black dotted lines in Fig. 4), as shown by the relative increase of decomposers over phytoplankton (orange versus blue and green lines in Fig. 4). Because decomposers have higher N:C and P:C ratios compared to phytoplankton, higher heterotrophy will tend to increase nutrients relatively to carbon in the seston (yellow areas in Fig. S5).

Whether fixers and non-fixers phytoplankton groups increase or decrease in density in Fig. 3 depends on the balance between two processes which regulate the availability of nutrients for phytoplankton: *(i)* the net mineralization of nutrients, *m*, that is a fixed parameter, and *(ii)* nutrient uptake by decomposers, *ϕ_I_*_,*P*_ and *ϕ_I_*_,*N*_, which are modulated by the stoichiometry of lake detritus (*α_D_*, *β_D_*) and their match to decomposers stoichiometry (*α_B_*, *β_B_*)

**Figure 3:**
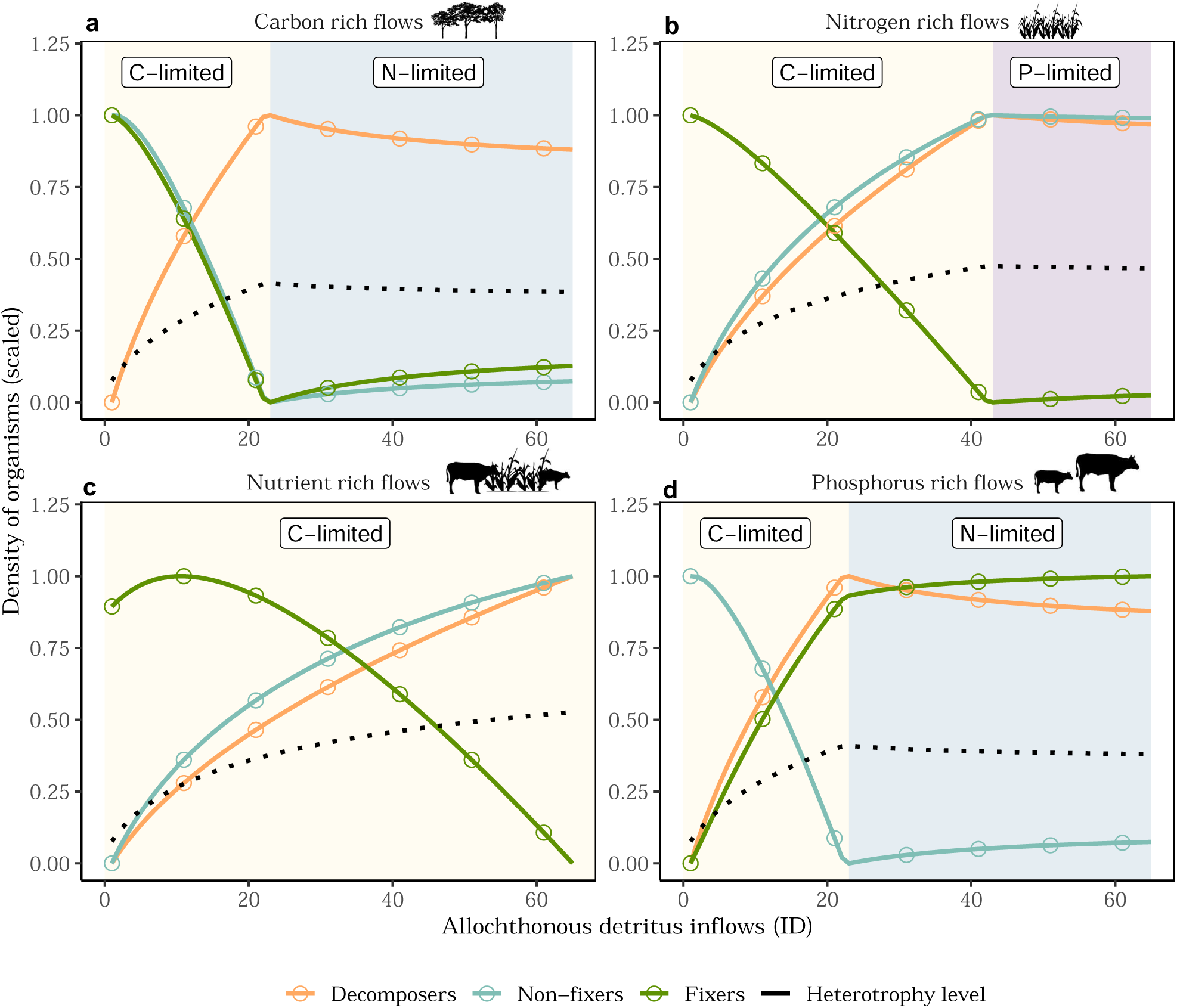
Allochthonous inflows promote heterotrophy until change in decomposers’ limitation. Densities of decomposers, fixers and non-fixers (in carbon unit), and heterotrophy level (black dotted lines; ratio of density of decomposers divided by the summed densities of phytoplankton and decomposers) along the gradient of allochthonous inflows (*I_D_*). Values are scaled since we are only interested in the change along the gradient of allochthonous inflows and not the net values. The panels (a-d) correspond to the different scenarios of stoichiometry of the allochthonous inflows (see Methods and Appendix A for parameter values). Blue and purple areas indicate regions where the decomposers are limited by nitrogen or phosphorus respectively (as compared to carbon limitation in yellow). The same figure but unscaled is available in Fig. S6.

**Figure 4:**
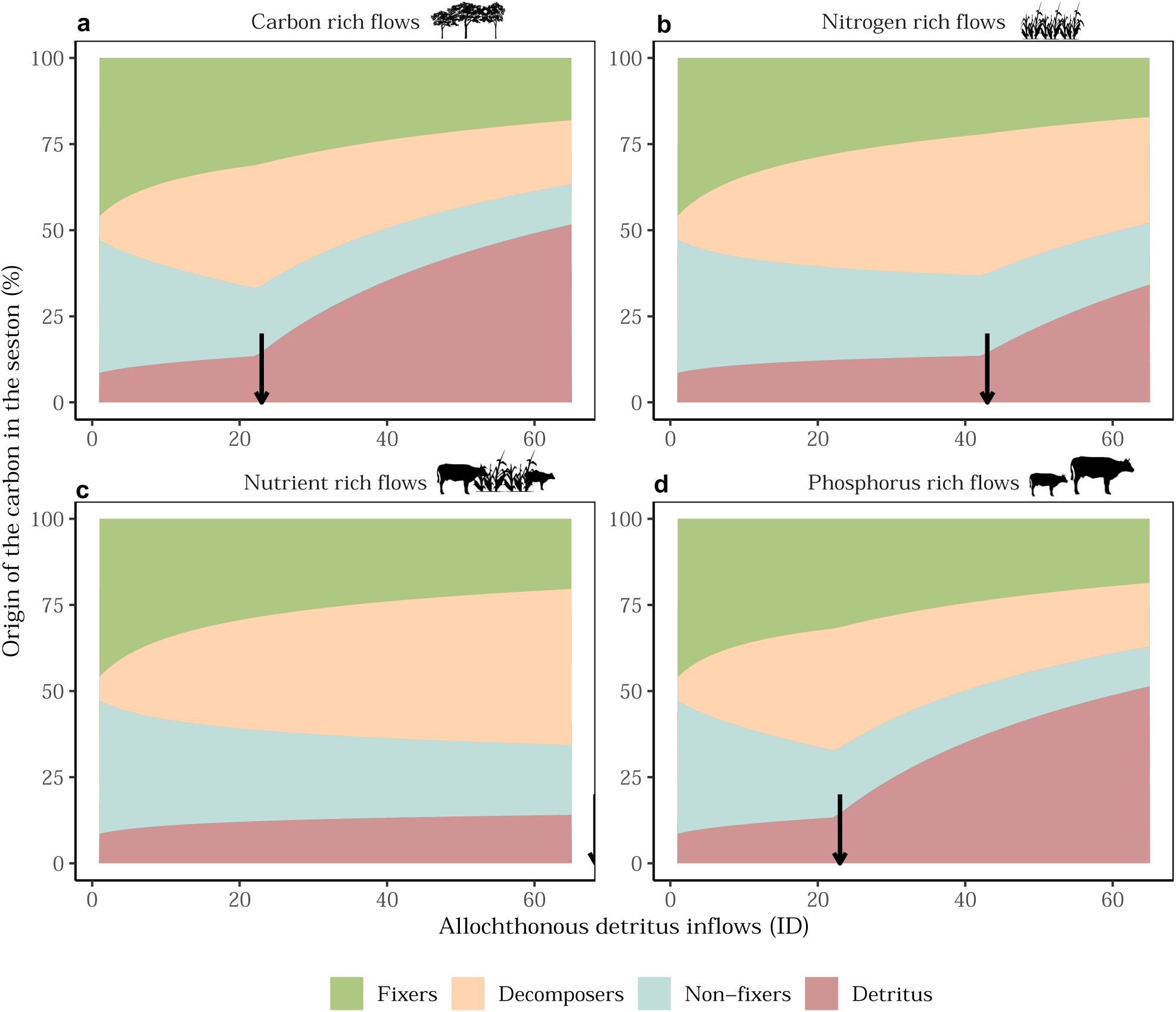
Partition of carbon in the seston from the different functional groups of organisms and the detritus. Composition of the carbon in the seston decomposed by the different functional groups of organisms and the detritus. The panels (a-d) correspond to the different scenarios of stoichiometry of the allochthonous inflows (see Methods and Appendix A for parameter values). The arrows indicate when there is a change in the limitation of decomposers from carbon to nutrients (either nitrogen or phosphorus). When allochthonous inflows alleviate decomposer carbon limitation (arrow), there is an abrupt change in the ecosystem’ ability to regulate allochthonous inflows, from which the sestonic elemental ratios become increasingly determined by the allochthonous inflows.

When allochthonous inflows are carbon-rich or nitrogen-rich, allochthonous inflows lower the P:C ratio of detritus because *β_A_ < β_D_* (Fig. S2). Since the stoichiometric mismatch between detritus and decomposers for phosphorus is already in the direction of detritus poorer in P than needed (*β_D_ < β_B_*), the mismatch increases (*i.e.,* |*β_B_* − *β_D_*| increases; Fig. S3). Consequently, decomposers need to immobilize more phosphorus to maintain their homeostasis (*i.e., ϕ_P_*increases). As such, competition from decomposers to fixers increases for phosphorus availability as emphasized by the decreasing strength of the indirect facilitation of decomposers on fixers (orange line in Fig. 5a-b), as well as the decrease of fixer population density (Fig. 3a-b).

**Figure 5:**
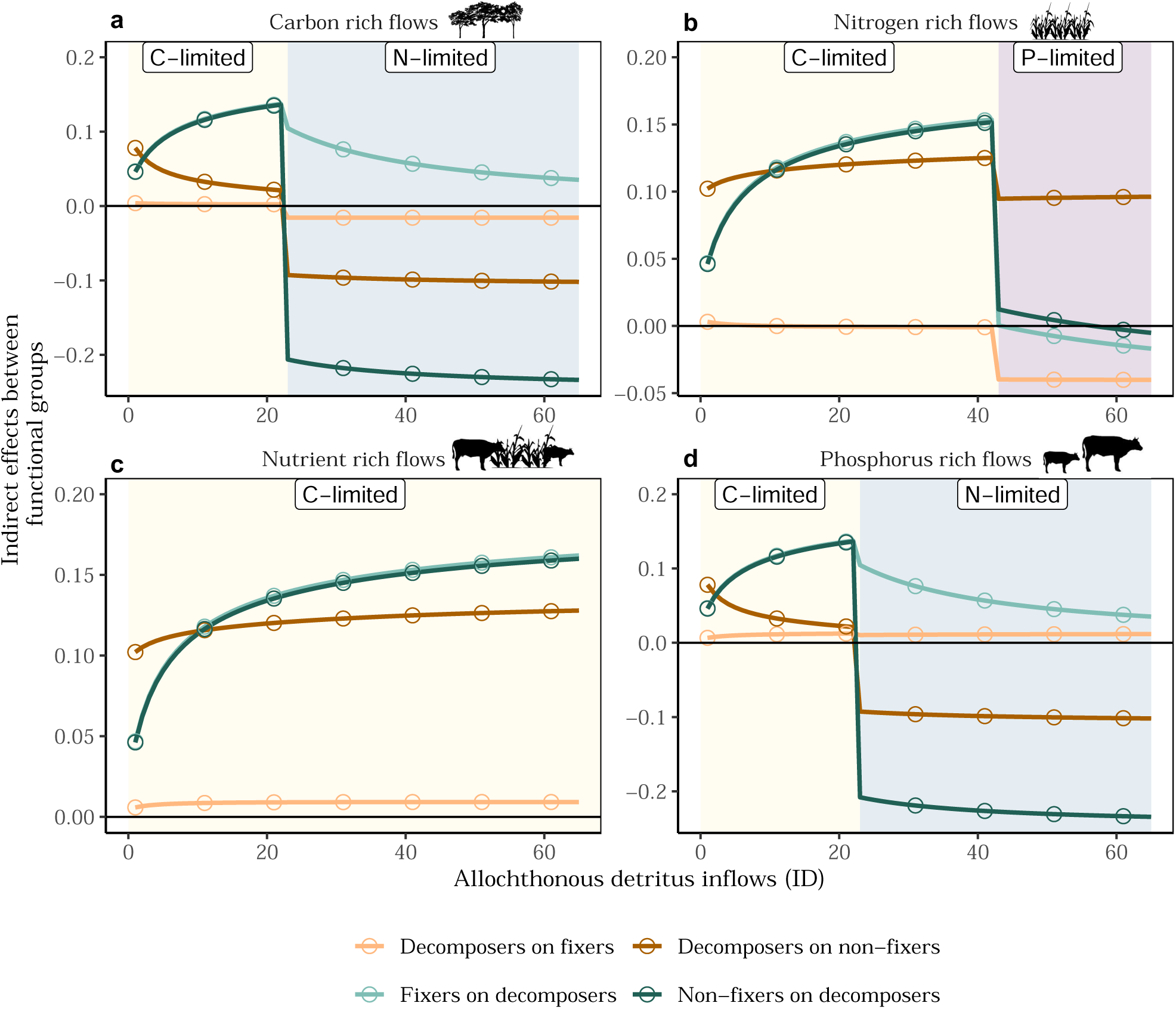
Indirect effects between decomposers and fixers or non-fixers along the gradient of allochthonous inflows (***I_D_***). Indirect effects between decomposers and fixers or non-fixers along the gradient of allochthonous inflows (*I_D_*). When allochthonous inflows alleviate decomposer carbon limitation, there is an abrupt change from indirect facilitation to indirect competition dominating the system due to shared limited resources between functional groups. The panels (a-d) correspond to the different scenarios of stoichiometry of the allochthonous inflows (see Methods and Appendix A for parameter values). Note that the indirect effects of decomposers on fixers become less (*resp.* more) positive for the carbon-rich and nitrogen-rich scenarios (*resp.* phosphorus and nutrient-rich scenarios) under moderate allochthonous inflows (*I_D_*). Blue and purple areas indicate regions where the decomposers are limited by nitrogen or phosphorus respectively (as compared to carbon limitation). Negative values indicate exploitative competition while positive values correspond to indirect facilitation.

For similar reasons, the N:C ratio of allochthonous inflows and the mismatch between detritus and decomposers for nitrogen control the dynamics of the non-fixer phytoplankton along the gradient of allochthonous inflow (Appendix A).

### High allochthonous inflows deregulate the control of seston stoichiometry

When subsidies are not enriched both in N and P, large amounts of spatial flows alleviate decomposers’ carbon limitation, and decomposers switch to nitrogen or phosphorus limitation (blue or purple areas in Fig. 3). This leads to drastic changes in the dynamics and functioning of the aquatic ecosystem, and the indirect effects between functional groups that become mainly dominated by competition (*e.g.,* see the differences between the nutrient-rich scenario and the other scenarios in Fig. 5). For example, high C-rich and P-rich inflows switch the carbon limitation of decomposers to nitrogen limitation, inducing a strong exploitative competition for nitrogen between non-fixers and decomposers. By contrast, high N-rich inflows trigger a switch from carbon to phosphorus limitation leading to strong competition between fixers and decomposers (Fig. 5). Dynamics are in this case increasingly driven by competition, which leads to the decrease in decomposer density and the increase of the phytoplankton: the ecosystem is thereby pushed back to slightly higher autotrophic functioning relative to heterotrophy (Fig. 3).

### High allochthonous inflows hinder stoichiometric homeostasis

Lastly, this change in decomposers’ limitation has a consequence for the stoichiometry of the lake (Fig. 4). When nutrient-limited, decomposers are less able to regulate the sustained increase of allochthonous carbon by decomposition, leading to a remarkable increase of the fraction of the seston occupied by detritus (see after arrows in Fig. 4). This loss of ability to regulate allochthonous inflows of detritus makes the seston stoichiometry increasingly reflecting stoichiometry of allochthonous inflows, which corresponds to a decrease in ecosystem’s homeostasis (see change in slopes between N- and P-compared to C-limited decomposers in Fig. 6a-b). With the loss of stoichiometric homeostasis, the elemental ratios of the seston become primarily driven by the stoichiometry of allochthonous inflows, which brings the ecosystem closer or father away from the Redfield ratio depending on its initial deviation in absence of allochthonous inflows and the stoichiometry of allochthonous inflows (Fig. S5). For example, when inflows are carbon rich, change in decomposers’ limitation from carbon to nitrogen, lowers the N:C ratios of the seston toward the Redfield ratio, and slows-down the increase of P:C in the seston (Fig. S5).

**Figure 6:**
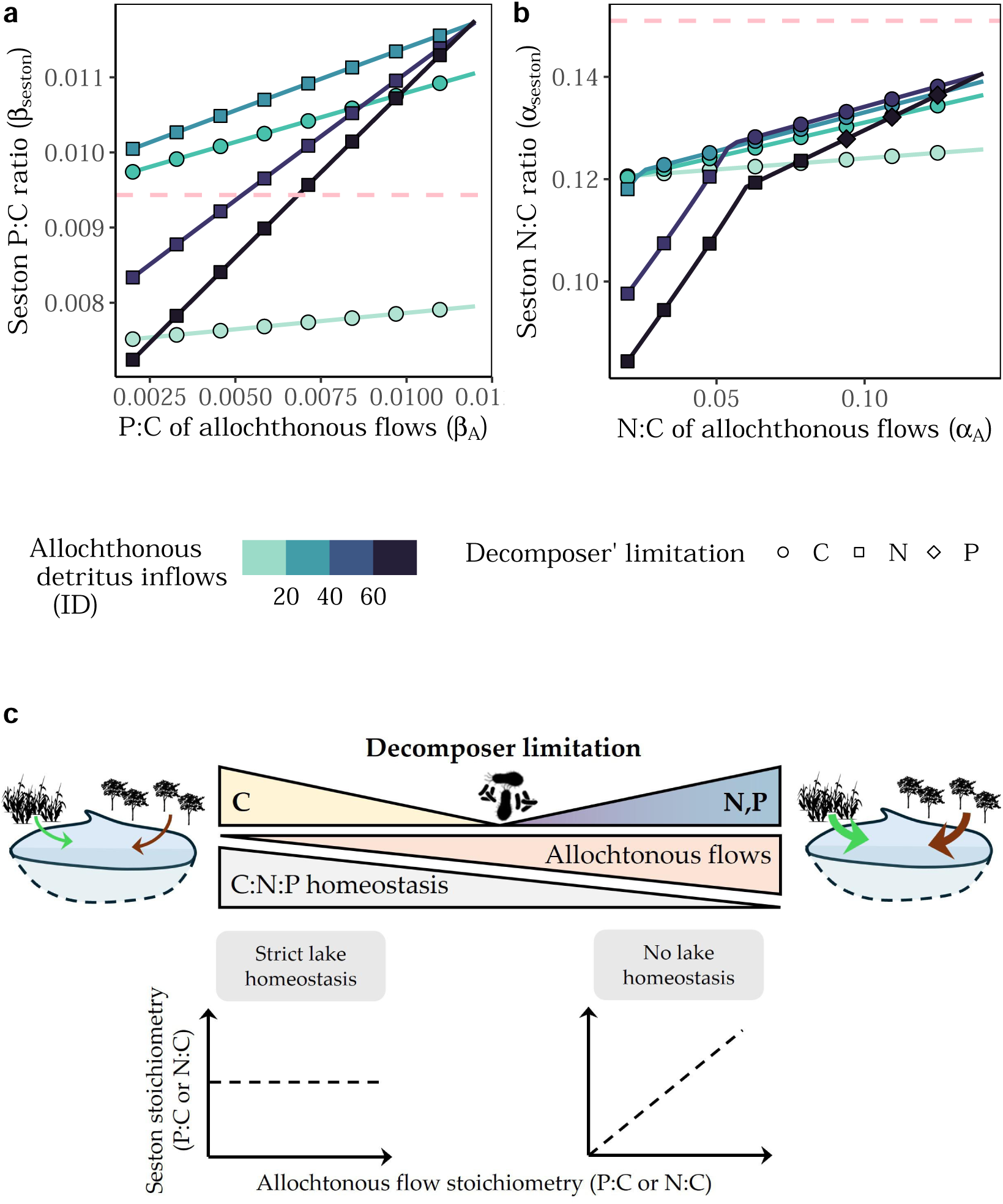
Homeostasis of sestonic C:N:P ratios. (Stoichiometric homeostasis at the ecosystem extent refers to the ability of the seston to maintain fixed elemental ratios (y-axis) despite stoichiometric changes of allochthonous inflows (x-axis). The homeostasis decreases non-linearly with the magnitude of allochthonous inflows (colour gradient). Yet, when decomposers carbon’ limitation is alleviated by the allochthonous flows, homeostasis is further reduced due to lower ability of decomposers to regulate allochthonous inflows (change from circle to square or diamond points in panel b). When decomposers are nutrient limited, allochthonous flows determine the sestonic elemental ratio and can make it deviate from or converge to the Redfield ratio (dashed pink line). (c) Illustrative interpretation of our results suggesting a gradual loss of lake’ stoichiometric homeostasis as a result of the allochthonous flows and the changes in decomposers’ limitation (yel_2_lo_4_w-blue/purple gradient).

## Discussion

We analyzed a stoichiometric model of aquatic ecosystem dynamics and our results emphasize the direct and indirect (through community dynamics) mechanistic pathways through which the quantity and quality of allochthonous inflows affect the stoichiometry of lakes. High levels of detrital allochthonous inflows can destabilize ecosystem’ functioning by alleviating decomposer carbon limitation, which promotes competitive interactions and the loss of ecosystem’ ability to regulate carbon inflow (*i.e.,* its stoichiometric homeostasis; Fig. 6c) by a shift of decomposers limitation from carbon to nutrient. We discuss these results in light of the Redfield ratio and provide perspectives for the functioning of aquatic ecosystems and meta-ecosystem ecology.

### Moderate allochthonous inflows promote heterotrophy and modulate decomposer-phytoplankton interactions

When allochthonous inflows are moderate such that decomposers remain carbon-limited, the aquatic ecosystem is first pushed toward higher heterotrophic functioning because of higher decomposition over autotrophic production from phytoplankton. These results are consistent with a rich body of literature emphasizing that bacteria production and decomposition in lakes can strongly rely on allochthonous inflows (Jonsson et al., 2001; Kritzberg et al., 2004, 2006), such that increase in allochthonous inflows can increase the ratio of heterotrophic to autotrophic production (*i.e.,* heterotrophy; Jansson et al., 2000; Bergströ m & Jansson, 2000). Yet, by quantifying indirect effects between functional groups, we find that allochthonous inflows mainly promote indirect facilitation when decomposers remain carbon-limited. Such facilitative interactions are due to the increasing mineralization of nutrients by decomposers and the production of detritus by phytoplankton, which could extend decomposers’ niche when allochthonous inflows are too small for decomposers to persist alone in the ecosystem. In such coupled brown-green ecosystems where decomposition and primary production coexist (Zou et al., 2016), experiments and theory have suggested that decomposers and primary producers positively or negatively interact depending on their limitation and their stoichiometric constraints (Daufresne & Loreau, 2001; Danger et al., 2007a). As such, we find that changes in decomposer density lead to cascading responses of phytoplankton density. In our model, the key mechanism underlying the effect of allochthonous inflows on the interaction between decomposers and producers is the mismatch between the demand in elements of decomposers and the elemental compositions of their detritus: *β_B_* − *β_D_* for phosphorus and *α_B_* − *α_D_* for nitrogen. Specifically, when allochthonous inflows increase the stoichiometric mismatch, they promote more competitive interactions between phytoplankton and decomposers due to higher immobilization of nutrients by decomposers, and it therefore lowers the density of phytoplankton.

In that sense, our results suggest that the quality of the allochthonous inflows determine whether allochthonous inflows act as a subsidy for phytoplankton (*i.e.,* positively fostering phytoplankton growth; *sensu* Polis et al., 1997) or as an inhibitory resource that limit their growth (Brett et al., 2009; Kelly et al., 2014). In other contexts, such stoichiometric mismatch between decomposers and their detritus has been shown to control the indirect effect of herbivores on nutrient availability for plants (Cherif & Loreau, 2013), and to reduce assimilation efficiency of consumers (Anderson et al., 2005; Hillebrand et al., 2009). At larger spatial scales, stoichiometric mismatch between organisms and their resources determine the sign of the emergent spatial feedback between aquatic and terrestrial ecosystems and the effects of allochthonous inflows on landscape functioning (Pichon et al., 2023). This stoichiometric constraint on growth may therefore add-up to the physical effect of allochthonous inflows on light availability, and further reinforce the level of heterotrophy in lakes and streams (Kelly et al., 2014; Dutton et al., 2018).

### Quality of allochthonous inflows on ecosystem functioning

Our model stresses three different effects of the quality of spatial resource flows, which determines *i)* the density and limitation of decomposers, *ii)* the relative density of phytoplankton groups and their reciprocal indirect effects, and *iii)* the elemental ratios in the seston. Our results extend previous findings on sestonic elemental ratios (*e.g.,* Hessen et al., 2003) by showing they emerge from the interaction between the stoichiometry of allochthonous inflows, the limitation of decomposers, and the level of lake heterotrophy. In our model, when decomposers are carbon-limited, they can regulate detritus inflow. Variations in seston elemental ratios are therefore driven by the changes in the relative density of decomposers compared to phytoplankton based on differences in elemental ratios between both functional groups.

Yet, when detritus inflows were high such that decomposers become nutrient-limited, detritus inflows are more weakly regulated and seston elemental ratios are increasingly determined by the stoichiometry of allochthonous inflows. Previous modelling approaches on the stability of N:P ratio in aquatic ecosystems, and particularly oceans, mostly considered the competition between fixers and non-fixers as the underlying homeostasis mechanism of the elemental ratios (Tyrrell, 1999). Yet, such models could not explain the strong deviations from the Redfield ratio in some regions (Martiny et al., 2013), as well as the relative enrichment in carbon of small lakes or coastal areas as compared to off-shore areas (Sterner et al., 2008). Our results shed light on these deviations by providing a mechanistic understanding of the homeostasis of sestonic elemental ratios driven by allochthonous inflows, interactions between decomposers and phytoplankton, and their respective stoichiometric constrains.

Empirical observations in various lakes (Box 1; Hessen et al., 2009; Prater et al., 2017; Soranno et al., 2019) have emphasized the importance of the regional context to understand lake dynamics and the stoichiometry of the seston in the lakes. Croplands for instance have been shown to increase the N:P ratio of lakes in watersheds (Arbuckle & Downing, 2001; Collins et al., 2017), and high riparian vegetation cover has been shown to increase organic carbon in lakes (Larsen et al., 2011). This has also been shown in recent experimental work where different qualities of additions in the experimental lakes led to contrasted effects on seston elemental ratios (Calderó-Pascual et al., 2022). In the seminal study by Sterner et al. (2008), small lakes were found to have larger deviations from the Redfield ratio. Our study allows providing a mechanistic basis for those observations since small lakes are more sensitive to large allochthonous inflows, and might therefore be less able to regulate allochthonous inflows.

### Perspectives for model extensions

In our model we made some hypotheses that we would like to discuss here. First, we assumed that allochthonous and autochthonous detritus were equally preferred by decomposers since only one compartment of detritus was considered. Yet, experiments suggest that despite autochthonous (local) resources might not always be sufficient to support decomposers growth, autochthonous resources are generally preferred over allochthonous ones (Kritzberg et al., 2004, although see Attermeyer et al., 2014). Moreover, under different qualities of allochthonous inflows, bacteria can adjust their growth efficiency and further promote their own dominance in humic lakes (Roiha et al., 2016). Moreover, as phytoplankton that can adjust their stoichiometry to that of their resource (Sterner & Elser, 2002), competitive exclusion in decomposer communities can lead to an overall adjustment of their stoichiometry to that of the detritus (Danger et al., 2008). This stoichiometric plasticity is an overlooked component of the homeostasis of the seston in aquatic ecosystems. Our modelling results provide in this sense a minimalistic picture to understand the complex interactions between phytoplankton and decomposers in a spatial context. Nevertheless, dissecting the indirect interactions between functional groups illustrates how bridging the gap between community ecology and ecological stoichiometry provides a more mechanistic understanding of aquatic ecosystem dynamics.

Secondly, one key factor we did not account for is the interactive effect between allochthonous inflows and light availability in these humic lakes or streams, since inflows further decrease primary production by decreasing light availability, independently of the quality of the detritus flows. Models explicitly accounting for light availability as a resource show that light attenuation can highly structure community composition along the water column (Jäger & Diehl, 2014), which can cascade-up to higher trophic levels in lakes (Kelly et al., 2014). We expect that accounting for light availability in the present bacteria-phytoplankton model would further increase the positive effect of allochthonous inflows on heterotrophic production by decreasing light availability, thereby hindering photosynthesis (Ask et al., 2009; Thrane et al., 2014).

### Deviations from the Redfield ratio in lakes

Open oceans have small deviations from the Redfield ratio (Sterner et al., 2008), which might be attributed to the composition of planktonic communities or species-specific constraints (Klausmeier et al., 2004; Weber & Deutsch, 2010). Allochthonous inflows have negligible effect on oceans’ stoichiometry due to the dilution effect. By contrast, lakes are open to large inflows from neighboring ecosystems, particularly small lakes which dilution effect likely explain their large deviations from the Redfield ratio observed (Sterner et al., 2008). Our modeling study allows dissecting the mechanisms underlying these deviations. We show that allochthonous inflows can promote decomposition over primary production and can induce a switch in the limitation of decomposers. Together theses processes push the ecosystem further away from the Redfield ratio. In aquatic ecosystems with low primary production that are highly dependent on allochthonous inflows such as lakes with high riparian cover (Gounand et al., 2018), allochthonous inflows might contribute largely to the stoichiometry of the seston and to their deviations from the Redfield ratio.

Our results therefore suggest a dynamical view of the Redfield ratio of ecosystems in space and in time. Across landscapes, differences in the quantity and the stoichiometry of allochthonous inflows due to catchment properties or the type of neighboring ecosystems –nutrient-rich inflows from agricultural areas *versus* carbon-rich litter from forest (Sitters et al., 2015; Collins et al., 2017; Prater et al., 2017)– might generate a mosaic of aquatic ecosystems across the landscape with heterogeneous seston’ stoichiometry (Collins et al., 2017; Soranno et al., 2019). Over time, deviations from the Redfield ratio in lakes might constantly change depending on the quantities and stoichiometry of resources exported from neighboring terrestrial ecosystems (*e.g.,* higher during growing season (Gudasz et al., 2012) or during organism migration (Subalusky et al., 2017; Peller et al., 2023), see Appendix D for a model extension over time). By bridging stoichiometry and community dynamics in aquatic ecosystems, our study highlights mechanisms of homeostasis and the cascading effects of allochthonous inflows on community composition, indirect effects, functioning, and sestonic elemental ratios.

## Conflict of interest disclosure

The authors of this article declare that they have no financial conflict of interest with the content of this article.

## Author contribution

B.P, T.D, I.G designed the study with inputs from F.G. B.P performed research and wrote the paper. All authors contributed to the subsequent versions.

## Funding

B.P acknowledges a doctoral fellowship from the chaire Modélisation Mathématique et Biodiversité of VEOLIA-Ecole Polytechnique-MNHN

## Notes

### Competing Interest Statement

The authors have declared no competing interest.

