## Supplementary file for "Do allochthonous flows explain deviations from the Redfield ratio in lakes?"

### Supplementary Material for
### Do allochthonous flows explain deviations from the Redfield ratio in lakes?

#### Contents

**Appendix S1: Supplementary methods and results** **S1**

**Appendix S2: Supplementary figures and tables** **S9**

**Appendix S3: Two-species model** **S16**

**Appendix S4: Time-varying stoichiometry of allochthonous flows** **S26**

### Appendix S1: Supplementary methods and results

#### *Supplementary results on fixers dynamics*

For similar reasons, the N:C ratio of allochthonous inflows and the mismatch between detritus and decomposers for nitrogen ($|\alpha_B - \alpha_D|$; Fig. S3), control the dynamics of the non-fixer phytoplankton along the gradient of allochthonous inflow (blue lines in Fig. 3a-b). With nitrogen-rich allochthonous inflows, decomposers immobilize less nitrogen, which promotes stronger mutual indirect facilitation between non-fixers and decomposers (dark green and brown lines in Fig. 5b-c), and increase non-fixer density (blue lines in Fig. 3b-c) before decomposers change from carbon to nutrient limitation.

It is noteworthy that under very low allochthonous inflows the presence of phytoplankton is a necessary condition for decomposers to persist (*i.e.*, orange line for $I_D = 0$ in Fig. S4). By producing detritus, phytoplankton allows decomposer population to increase, therefore explaining the positive effect of phytoplankton on decomposers (light and dark blue lines in Fig. 5a-d).

#### *Model and parameters*

*Derivation of $\phi_{I,N}$, $\phi_{I,P}$, $\phi_D$*

Stoichiometric homeostasis of decomposers implies:

$$\frac{dB_N}{dt} = \alpha_B \frac{dB_C}{dt} = \frac{\alpha_B}{\beta_B} \frac{dB_P}{dt}$$

which constrains the immobilization and decomposition flows :

$$\phi_{I,N} = \frac{\alpha_B - \alpha_D}{\alpha_B}\phi_D \text{ , with } \alpha_D = \frac{D_N}{D_C}$$
$$\phi_{I,P} = \frac{\beta_B - \beta_D}{\beta_B}\phi_D \text{ , with } \beta_D = \frac{D_P}{D_C}$$
$$\phi_{I,N} = \frac{(\alpha_B - \alpha_D)\beta_B}{(\beta_B - \beta_D)\alpha_B}\phi_{I,P}$$

When decomposers are N- (*resp.* P-)limited, decomposers immobilize nitrogen (*resp.* phosphorus) and modulate the immobilization of $P$ (*resp.* $N$) to maintain their homeostasis. Mathematically speaking, $\phi_{I,N}$ constrains $\phi_D$ and $\phi_{I,P}$ when decomposers are N-limited while $\phi_{I,P}$ constrains $\phi_D$ and $\phi_{I,N}$ when decomposers are P-limited. Under such limitation, a decrease in N:C ratio of detritus limits the decomposition rate of decomposers (*i.e.*, $\phi_D$ decreases due to an increase of $\alpha_B - \alpha_D$). Similar reasoning can be done for phosphorus limitation with the $\phi_D$ decreasing with $\beta_B - \beta_D$). On the contrary, when decomposers are carbon limited, the decomposition process ($\phi_D$) constrains the uptake or release of nitrogen and phosphorus ($\phi_{I,N}, \phi_{I,P}$). This is summarized in the scheme below.

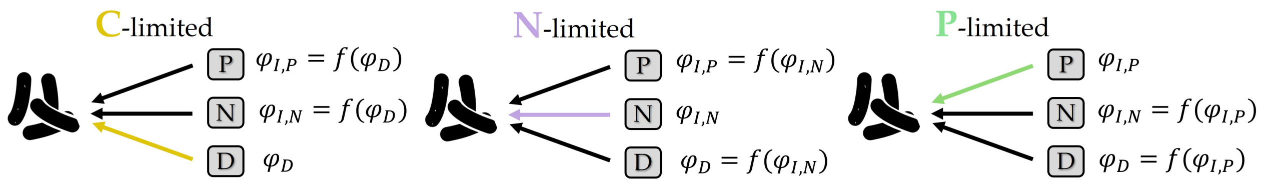


**Constrains upon immobilization and decomposition of nutrients and carbon from decomposers.**

Thus, we get the following formulas for the immobilization and decomposition flows for carbon and nitrogen-limited decomposers :

| | $\phi_{I,N}$ | $\phi_{I,P}$ | $\phi_D$ |
|---|---|---|---|
| **N-limited** | $a_N N B_C$ | $\frac{(\beta_B-\beta_D)\alpha_B}{(\alpha_B-\alpha_D)\beta_B}\phi_{I,N}$ | $\frac{\alpha_B}{\alpha_B-\alpha_D}\phi_{I,N}$ |
| **P-limited** | $\frac{(\alpha_B-\alpha_D)\beta_B}{(\beta_B-\beta_D)\alpha_B}\phi_{I,P}$ | $a_P P B_C$ | $\frac{\beta_B}{\beta_B-\beta_D}\phi_{I,P}$ |
| **C-limited** | $\frac{\alpha_B-\alpha_D}{\alpha_B}\phi_D$ | $\frac{\beta_B-\beta_D}{\beta_B}\phi_D$ | $e_B a_D D_C B_C$ |

The full expressions of $\phi_{I,N}, \phi_{I,P}, \phi_D$ are therefore given by:

$$\begin{cases} \phi_C = \min(\overbrace{e_B a_D D_C B_C}^{C-limitation}, \overbrace{\frac{\beta_B \phi_{I,P}}{\beta_B-\beta_D}}^{P-limitation}, \overbrace{\frac{\alpha_B \phi_{I,N}}{\alpha_B-\alpha_D}}^{N-limitation}) \\ \phi_{I,P} = \min(\overbrace{(\beta_B-\beta_D)\frac{\phi_D}{\beta_B}}^{C-limitation}, \overbrace{a_P P B_C}^{P-limitation}, \overbrace{\frac{(\beta_B-\beta_D)\alpha_B}{(\alpha_B-\alpha_D)\beta_B}\phi_{I,N}}^{N-limitation}) \\ \phi_{I,N} = \min(\overbrace{(\alpha_B-\alpha_D)\frac{\phi_D}{\alpha_B}}^{C-limitation}, \overbrace{\frac{(\alpha_B-\alpha_D)\beta_B}{(\beta_B-\beta_D)\alpha_B}\phi_{I,P}}^{P-limitation}, \overbrace{a_N N B_C}^{N-limitation}) \end{cases} \quad \text{(S1)}$$

*Assessing decomposer' limitation*

Practically speaking, we assessed the limitation of decomposers at system' equilibrium by quantifying the following limitation ratios:

$$\begin{aligned} S_{\text{C:N, lim}} &= \frac{(\alpha_B - \bar{\alpha}_D) a_B \bar{D}_C}{\alpha_B a_N \bar{N}}, \quad \bar{\alpha}_D = \frac{\bar{D}_N}{\bar{D}_C} \\ S_{\text{C:P, lim}} &= \frac{(\beta_B - \bar{\beta}_D) a_B \bar{D}_C}{\beta_B a_N \bar{N}}, \quad \bar{\beta}_D = \frac{\bar{D}_P}{\bar{D}_C} \\ S_{\text{N:P, lim}} &= \frac{a_N \bar{N}}{a_P \bar{P}}) \end{aligned}$$

, where the bar is used for equilibrium quantities.

In this case decomposer' limitations are:

- Carbon: when $S_{\text{C:N, lim}} < 1$ and $S_{\text{C:P, lim}} < 1$
- Nitrogen: when $S_{\text{C:N, lim}} < 1$ and $S_{\text{C:P, lim}} > 1$

- Phosphorus: when $S_{\text{C:N, lim}} > 1$ and $S_{\text{C:P, lim}} < 1$
- Nitrogen: when $S_{\text{C:N, lim}} > 1$, $S_{\text{C:P, lim}} > 1$ and $S_{\text{N:P, lim}} < 1$
- Phosphorus: when $S_{\text{C:N, lim}} > 1$, $S_{\text{C:P, lim}} > 1$ and $S_{\text{N:P, lim}} > 1$

*Assessing pairwise niche effects of functional groups*

We measured how the density of each functional group changed with the presence of others by measuring the change in functional group' density with and without others. This index, $NC_X, X \in \{F, O, B\}$, has been widely used in facilitation studies to quantify neighboring effects of a nurse species on a beneficiary one, and is given by (Díaz-Sierra et al., 2017):

$$NC_X = 2\frac{D_X - D_{\text{X only}}}{D_{\text{X only}} + |D_X - D_{\text{X only}}|} \tag{S2}$$

, where $D_X$ is the density of functional group $X$ with other functional groups, while $D_{\text{X only}}$ is the density of the functional group $X$ when alone in the ecosystem. This index is equal to -1 for cases of competitive exclusion and 2 for obligate facilitation (niche expansion).

*Parameter meaning*

*Parameter values*

We focused our analysis on the qualitative behavior that could emerge from the allochthonous inflow of nutrients and detritus in the aquatic ecosystem rather than exploring the full range of parameter values (that is virtually impossible).

Functional response parameters were taken from the literature and some of them were assumed to allow coexistence (all resource stocks and trophic levels having a positive

Table S1: **Parameters meaning and symbol.**

| Parameter | Unit | Meaning |
|---|---|---|
| **Abiotic flows** | | |
| $I_N$ | N.day$^{-1}$ | Nitrogen inflow in the aquatic ecosystem |
| $I_P$ | O.day$^{-1}$ | Phosphorus inflow in the aquatic ecosystem |
| $I_D$ | C.day$^{-1}$ | Detritus inflow in the aquatic ecosystem |
| $l_D$ | day$^{-1}$ | Loss rate of detritus in the aquatic ecosystem |
| $l_N$ | day$^{-1}$ | Loss rate of nitrogen in the aquatic ecosystem |
| $l_P$ | day$^{-1}$ | Loss rate of phosphorus in the aquatic ecosystem |
| $d_B$ | day$^{-1}$ | Loss rate of decomposers |
| $d_O$ | day$^{-1}$ | Loss rate of non-fixers |
| $d_F$ | day$^{-1}$ | Loss rate of fixers |
| **Biotic rates** | | |
| $m$ | day$^{-1}$ | Mineralization rate of decomposers |
| $e_B$ | dimensionless | Growth efficiency of decomposers |
| $a_N$ | day$^{-1}$ | Nitrogen consumption rate of decomposers |
| $a_P$ | day$^{-1}$ | Phosphorus consumption rate of decomposers |
| $a_D$ | day$^{-1}$ | Detritus consumption rate of decomposers |
| $\mu_F$ | day$^{-1}$ | Growth rate of fixers |
| $\mu_O$ | day$^{-1}$ | Growth rate of non-fixers |
| $s_B$ | day$^{-2}$ | Self-regulation rate of decomposers |
| $s_O$ | day$^{-2}$ | Self-regulation rate of non-fixers |
| $s_F$ | day$^{-2}$ | Self-regulation rate of fixers |
| **Stoichiometric ratio** | | |
| $\alpha_B$ | N:C (molar) | N:C ratio of decomposers |
| $\alpha_F$ | N:C (molar) | N:C ratio of fixers |
| $\alpha_O$ | N:C (molar) | N:C ratio of non-fixers |
| $\beta_B$ | P:C (molar) | P:C ratio of decomposers |
| $\beta_F$ | P:C (molar) | P:C ratio of fixers |
| $\beta_O$ | P:C (molar) | P:C ratio of non-fixers |

value) in each isolated ecosystem. This coexistence was also facilitated by the presence of the self-regulation term.

N:C ratios of organisms were constrained accounting for the lower N:C and P:C ratio of primary producers compared to decomposers (Elser et al., 2000; Cleveland & Liptzin, 2007; Buchkowski et al., 2019), and to the N:P ratio of 16 found in phytoplankton (Tyrrell, 1999). As such, we fixed the N:C ratio of decomposers to 0.15 but results were similar when $\alpha_B$ varied between 0.15 and 0.25 (Cleveland & Liptzin, 2007; Buchkowski et al., 2019).

Table S2: **Parameters values.**

| Parameter | Value |
|---|---|
| **Abiotic flows** | |
| $I_N$ | 5 |
| $I_P$ | 5 |
| $I_D$ | 1-100 |
| $l_D$ | 2 |
| $l_N$ | 1 |
| $l_P$ | 1 |
| $d_B$ | 0.1 |
| $d_O$ | 0.2 |
| $d_F$ | 0.2 |
| **Biotic rates** | |
| $m$ | 0.5 |
| $e_B$ | 0.5 |
| $a_N$ | 0.4 |
| $a_P$ | 0.4 |
| $a_D$ | 0.83 |
| $\mu_F$ | 0.24 |
| $\mu_O$ | 0.25 |
| $s_B$ | 0.1 |
| $s_O$ | 0.1 |
| $s_F$ | 0.1 |
| **Stoichiometric ratio** | |
| $\alpha_B$ | 0.15 |
| $\alpha_F$ | 0.125 |
| $\alpha_O$ | 0.125 |
| $\beta_B$ | 0.015 |
| $\beta_F$ | 1/128 |
| $\beta_O$ | 1/128 |

Table S3: **Parameters values for the quality of allochthonous inflows for the four different scenarios.**

| Parameter | Nutrient rich scenario | Phosphorus rich scenario | Nitrogen rich scenario | Carbon rich scenario |
|---|---|---|---|---|
| N:C allochthonous flows ($\alpha_A$) | 0.02 | 0.002 | 0.002 | 0.02 |
| P:C allochthonous flows ($\beta_A$) | 0.14 | 0.012 | 0.14 | 0.012 |

#### *Empirical data*

We aimed at providing a panorama of the elemental ratios of cross-ecosystem flows exported to aquatic ecosystems (ponds, lakes, and streams). For that, we gathered N:C, P:C and N:P ratios of resource allochthonous inflows. We started collecting elemental ratios from the studies used in the allochthonous inflow databases (Gounand et al., 2018; Pichon et al., 2023) or calculated them when two C, P or N flow estimates were provided in the same study. For each ratio, we also recorded the type of subsidies that is exported to aquatic ecosystems (*e.g.,* carcasses, mammal faeces, invertebrates) and the type of ecosystem from which those subsidies are exported (mostly grassland and forest). In total, we gathered 52 N:P ratios, 74 C:P ratios, and 123 C:N ratios of subsidies exported to aquatic ecosystems. We provide in the Zenodo repository the data associated with this database of elemental ratios of subsidies exported to aquatic ecosystems and the associated papers.

For Fig. 1d, we also represented the relative deviation of the elemental ratio of each subsidy to the Redfield ratio 116:16:1 for C:N:P using the following formula:

$$\text{Deviation (N:P)} = \frac{\text{N:P}_{\text{obs}} - \text{N:P}_{\text{Redfield}}}{\text{N:P}_{\text{Redfield}}}$$

, where $\text{N:P}_{\text{Redfield}} = 16$. The similar formula was applied to C:P and C:N with $\text{C:N}_{\text{Redfield}} = 116/16$ and $\text{C:P}_{\text{Redfield}} = 116$.

### APPENDIX S2: SUPPLEMENTARY FIGURES AND TABLES

Table S4: **Summary of elemental ratios of allochthonous inflows exported in lakes and streams.**
q25, q50 and q75 being the first, second (median) and third quantiles respectively. n is the number of data points.

| Ratio | n | min | q25 | q50 | q75 | max |
|---|---|---|---|---|---|---|
| N:P | 52 | 3.09 | 9.88 | 18.27 | 34.27 | 100.20 |
| C:P | 74 | 2.00 | 322.50 | 706.50 | 1132.97 | 7198 |
| C:N | 123 | 3.77 | 22.96 | 35.40 | 55.75 | 169.24 |

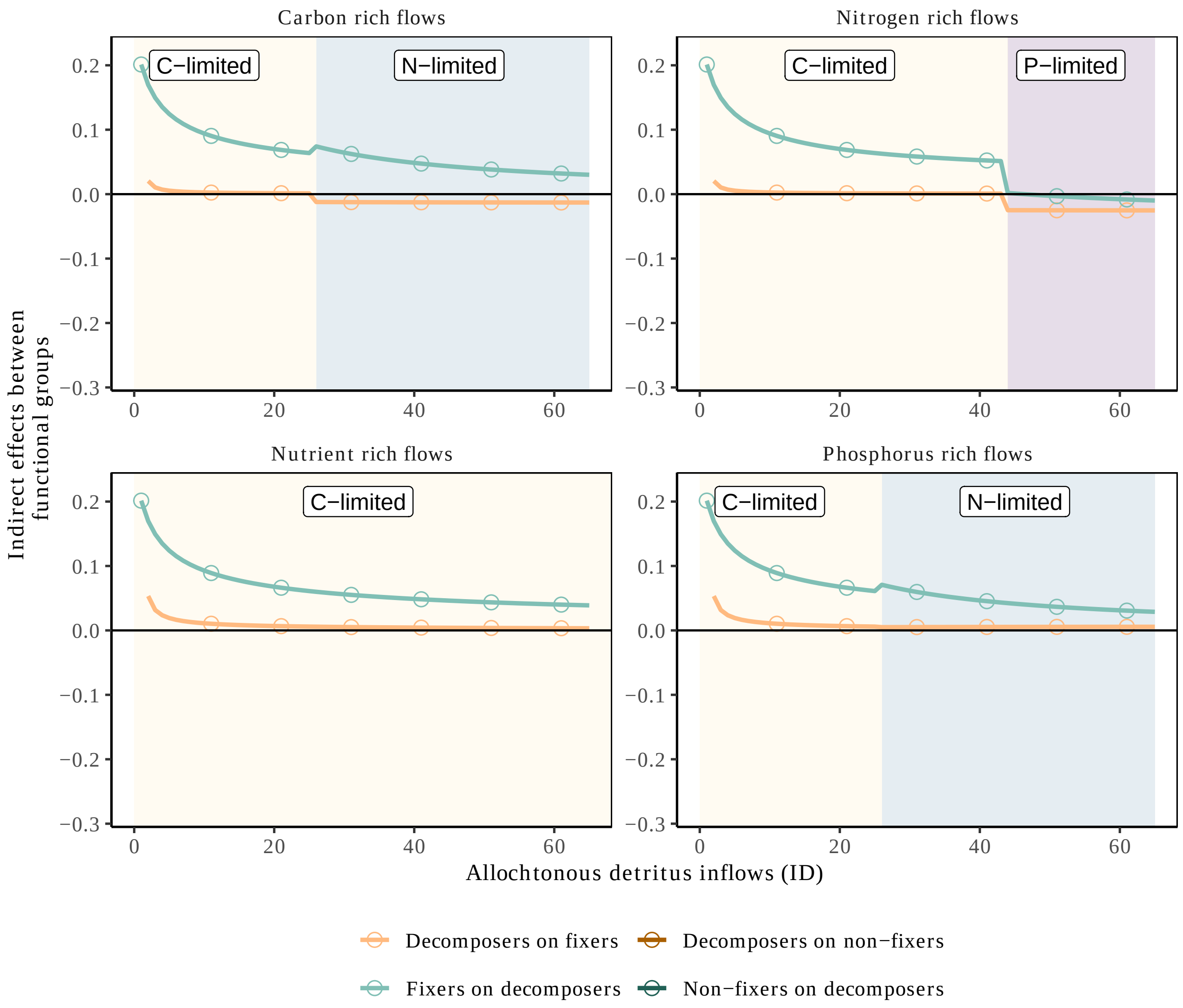


Figure S1: **Allochthonous inflows destabilize indirect effects between functional groups even when not controlling for the self-loop ($I_{ij} = s_{ij}$).**
Indirect effects between decomposers and fixers or non-fixers along the gradient of allochthonous inflows (ID). The panels correspond to the different scenarios of stoichiometry of the allochthonous flows (see Methods and Appendix A for parameter values). The blue and purple areas indicate regions where the decomposers are limited by nitrogen or phosphorus respectively (as compared to carbon limitation in yellow). Negative values indicate indirect competition while positive values correspond to indirect facilitation. There are no sign differences between this figure and Fig. 5.

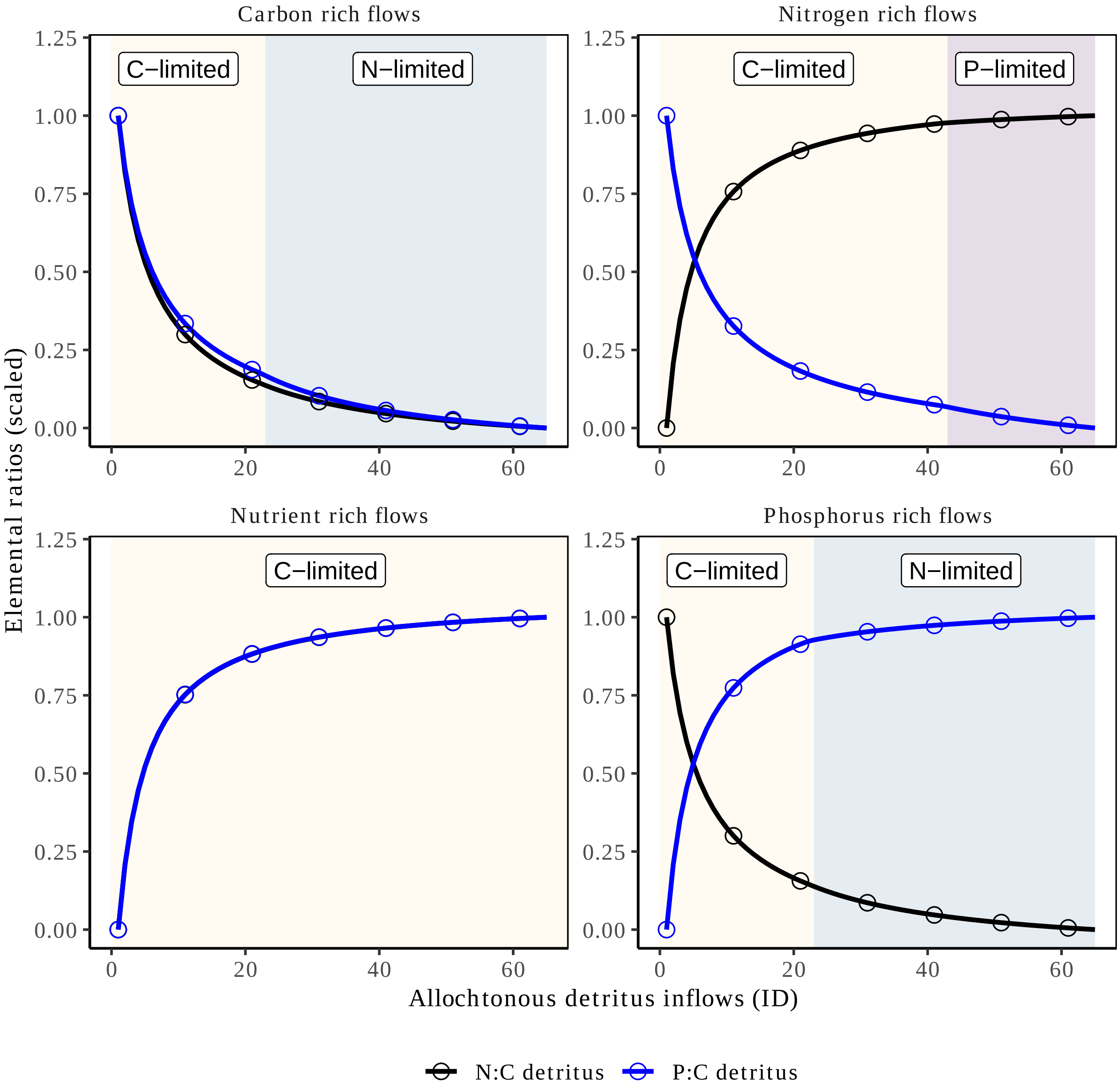


Figure S2: **Allochthonous inflows drive change in detritus stoichiometry.**
Change in N:C and P:C of detritus along the gradient of allochthonous inflows (ID). When flows are carbon rich and nutrient rich, the P:C ratio of detritus decreases non-linearly with increasing allochthonous inflows (and inversely for the two other scenarios). When flows are carbon rich and phosphorus rich, the N:C ratio of detritus decreases non-linearly with increasing allochthonous inflows (and inversely for the two other scenarios). The panels correspond to the different scenarios of stoichiometry of the allochthonous flows (see Methods and Appendix A for parameter values). The blue and purple areas indicate regions where the decomposers are limited by nitrogen or phosphorus respectively (as compared to carbon limitation in yellow).

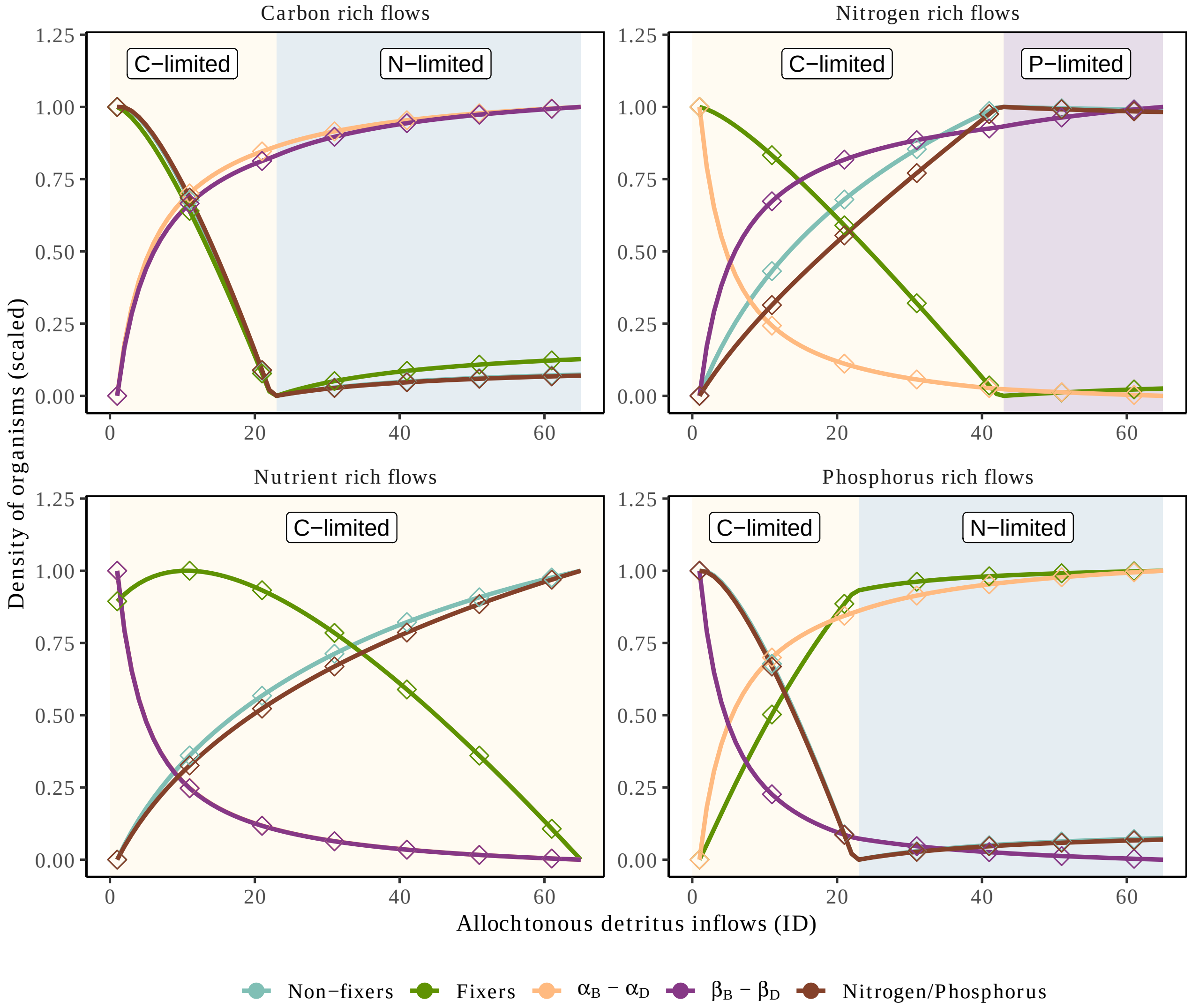


Figure S3: **Mechanisms of change in phytoplankton density along the gradient of allochthonous inflows.**
Change in phytoplankton (fixers and non-fixers) density and the stoichiometric mismatch ($\beta_B - \beta_D$ and $\alpha_B - \alpha_D$) between decomposers and their detritus along the gradient of allochthonous inflows (ID). Depending on the changes of $\beta_B - \beta_D$ along the allochthonous inflow gradient, fixers decrease (when $\beta_B - \beta_D$ increases) or increase in density (when $\beta_B - \beta_D$ decreases). This is similar for the changes of non-fixers but with $\alpha_B - \alpha_D$. The panels correspond to the different scenarios of stoichiometry of the allochthonous flows (see Methods and Appendix A for parameter values). The blue and purple areas indicate regions where the decomposers are limited by nitrogen or phosphorus respectively (as compared to carbon limitation in yellow).

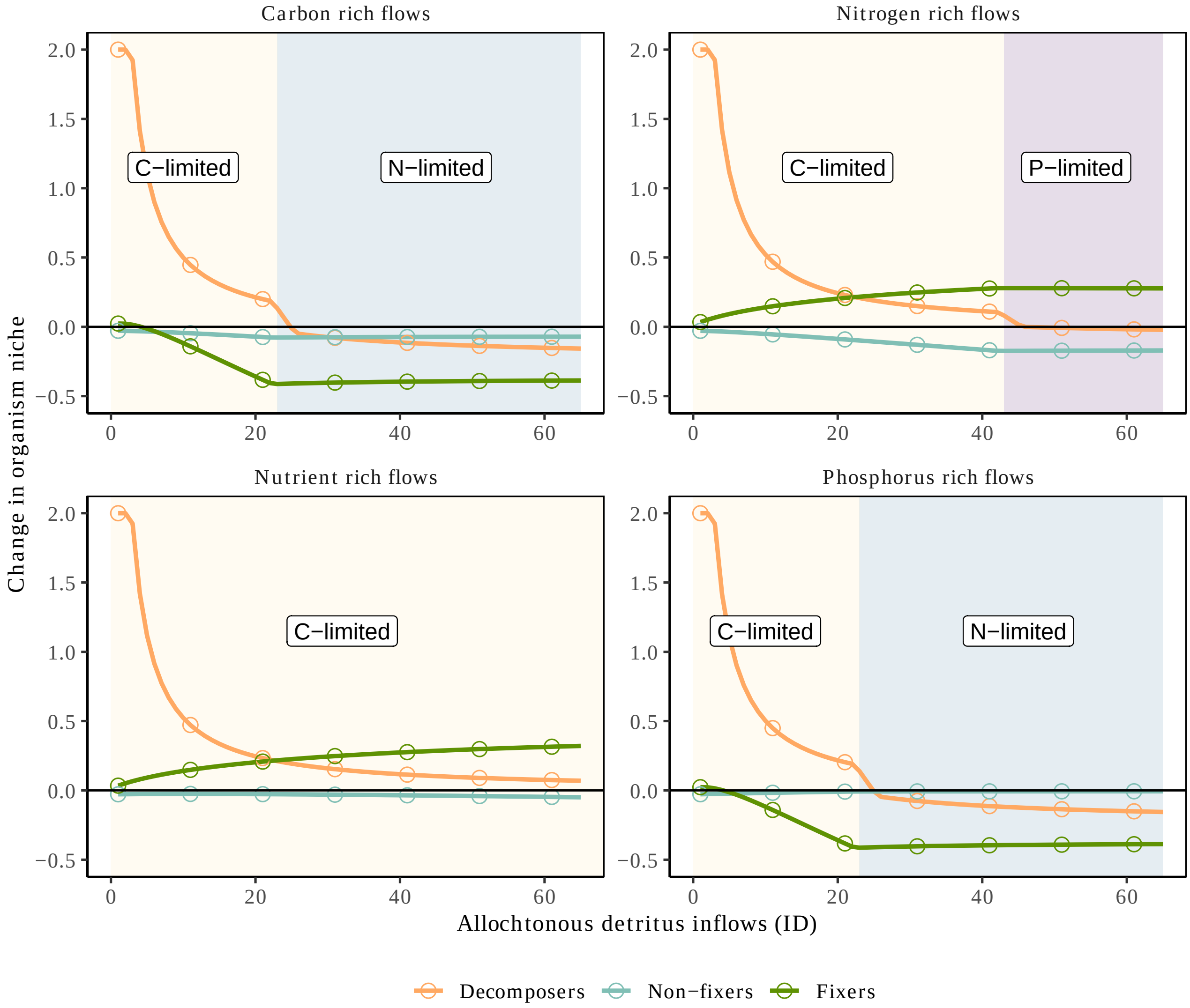


Figure S4: **Niche extension and contraction along the gradient of allochthonous inflows.** Change in the niche of each functional group of organism in the presence of all other functional groups (see Appendix A for the metric). Positive values indicate facilitation in the sense that the presence of other functional groups increases the density of the focal functional group. When values equal +2 it means that the focal functional group cannot persist without the other functional groups. By contrast, negative values indicate competition such that the presence of other functional groups decreases the density of the focal functional group. When values equal -1 it indicates competitive exclusion, which never occur in our model due to the self-regulation terms maintaining coexistence. The panels correspond to the different scenarios of stoichiometry of the allochthonous flows (see Methods and Appendix A for parameter values). The blue and purple areas indicate regions where the decomposers are limited by nitrogen or phosphorus respectively (as compared to carbon limitation in yellow).

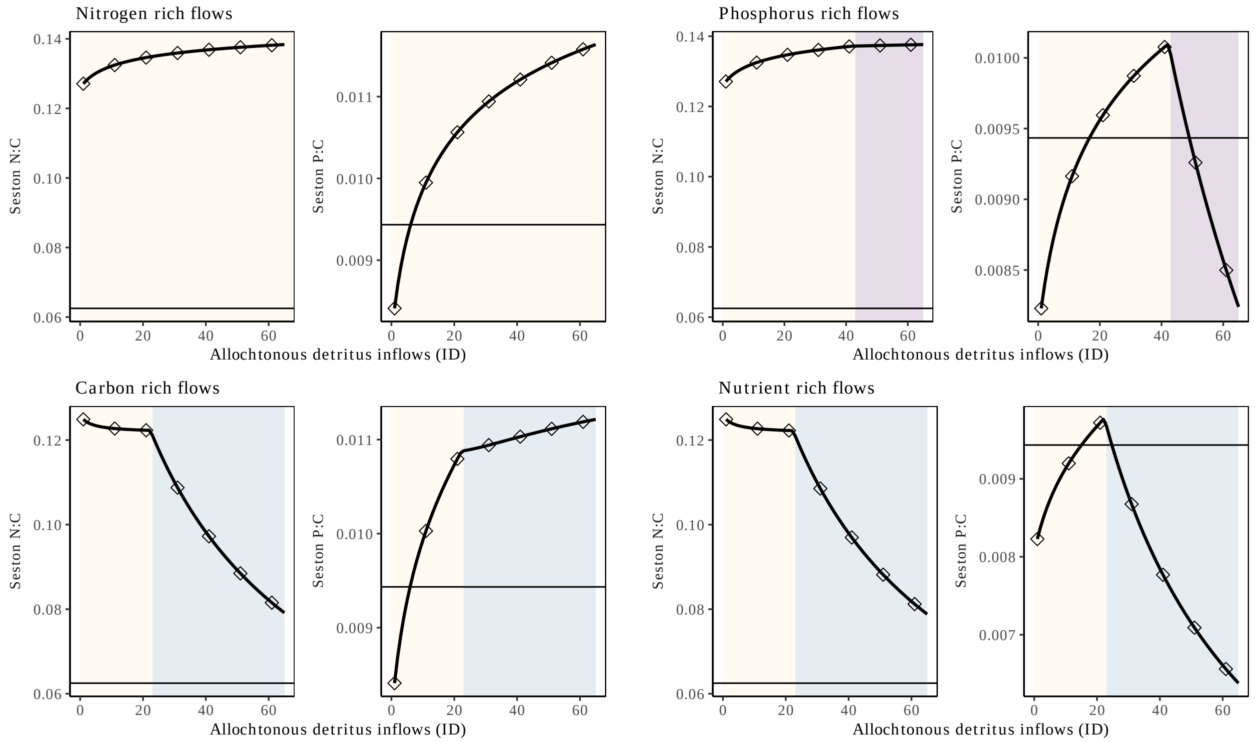


Figure S5: **Spatial stoichiometry of flows drive the stoichiometry in the seston.** Change of the N:C and P:C ratios in the seston along the gradient of allochthonous inflows (ID) for the different scenarios. We see that allochthonous inflows control the stoichiometry of the seston, especially when decomposers are N- or P-limited and cannot control and decompose allochthonous inflows as much as if decomposers were C-limited. The elemental ratios in the seston and their deviation from the Redfield ratio (black horizontal line) are therefore contingent on the stoichiometry of allochthonous inflows. The panels correspond to the different scenarios of stoichiometry of the allochthonous flows (see Methods and Appendix A for parameter values). The blue and purple areas indicate regions where the decomposers are limited by nitrogen or phosphorus respectively (as compared to carbon limitation in yellow).

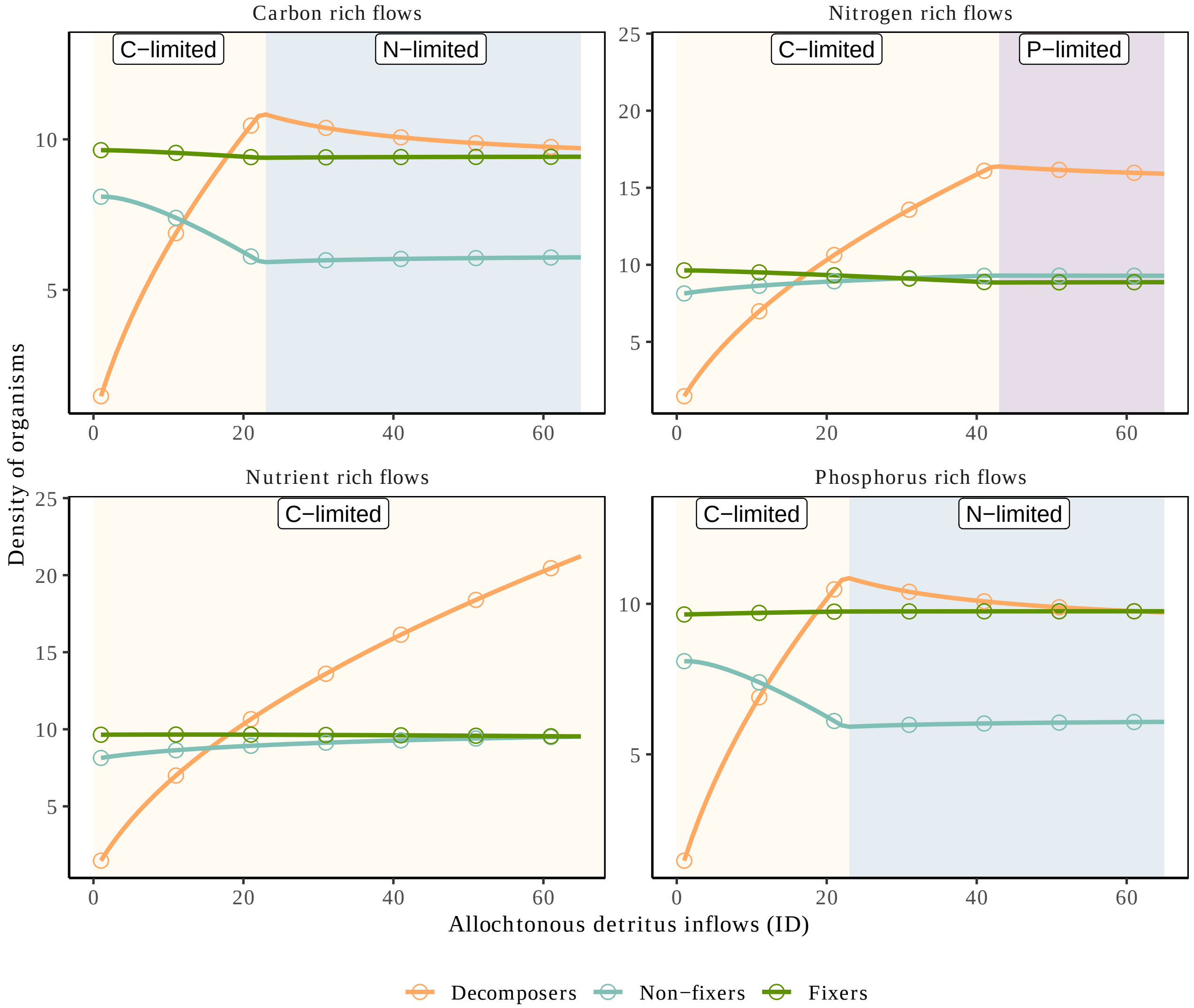


Figure S6: **Allochthonous inflows promotes heterotrophy until change in decomposer' limitation.**
Densities of decomposers, fixers and non-fixers (in carbon unit) along the gradient of allochthonous inflows ($I_D$). Compared to Fig. 3, densities are not scaled. The panels correspond to the different scenarios of stoichiometry of the allochthonous flows (see Methods and Appendix A for parameter values). The blue and purple areas indicate regions where the decomposers are limited by nitrogen or phosphorus respectively (as compared to carbon limitation in yellow).

### Appendix S3: Two-species model

In this section, we simplify the model to understand the two-species relationships. We focus on decomposers and fixers first and then investigate decomposers and non-fixers interaction.

#### *Decomposers and fixers*

Setting non-fixer phytoplankton population to zero ($O = 0$), we can study the dynamics the two-species model with only decomposers and fixers along a gradient of allochthonous inflows quantity and quality.

Increasing allochthonous inflows increases decomposers density. Whether fixers increase or not in density depends on the quality of allochthonous inflows (Fig. S7). Specifically when $\beta_B - \beta_D = \frac{B_P}{B_C} - \frac{D_P}{D_C}$ increases, this means that P:C of detritus decreases due to poor quality allochthonous inflows relatively to the detritus stoichiometry. Because detritus decrease in quality, decomposers need to take more and more from the phosphorus pool, which decreases fixers density whose dynamics are controlled by phosphorus availability. Consequently, decomposers only have a slight positive indirect effect on fixers (due to the balance between positive effect of mineralization and the negative ones from competition for phosphorus; Fig. S8). Fixers by contrast have a net facilitating effect on decomposers by the production of detritus. The dynamics change when decomposers switch from $C$ to $N$ or $P$ limitation (shaded area). In this case, decomposers compete more with fixers (mostly negative indirect effects), and symmetrically, has less and less indirect facilitating effects on decomposers (Fig. S8). Specifically, when decomposers are $P$-limited, the indirect effect of fixers on decomposers turn into a indirect competition since both share the same resources.

As explained in the main text, change in limitation of decomposers was also associated with loss of regulation of detritus inflows (Figs. S9,S10).

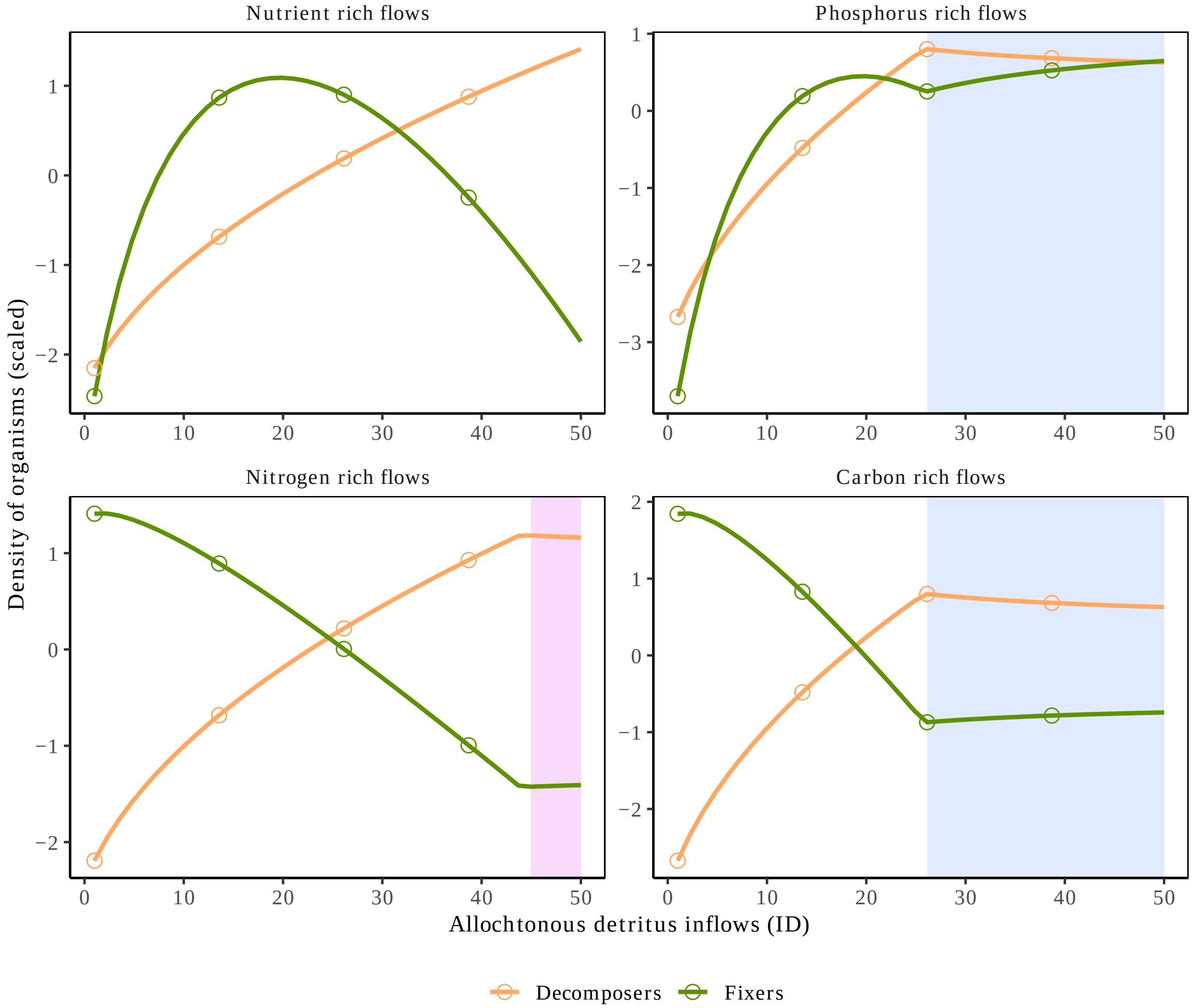


Figure S7: **Dynamics of fixers and decomposers along a gradient of allochthonous inflows with varying quality (facets).**

Scaled densities of decomposers and fixers (in carbon unit) along the gradient of allochthonous inflows (ID). The panels correspond to different stoichiometry of the allochthonous flows (low or high P:C and N:C). The blue and purple areas indicate regions where the decomposers are limited by nitrogen or phosphorus respectively (as compared to carbon limitation). The densities are scaled to allow comparison of the dynamics along the gradient of ID.

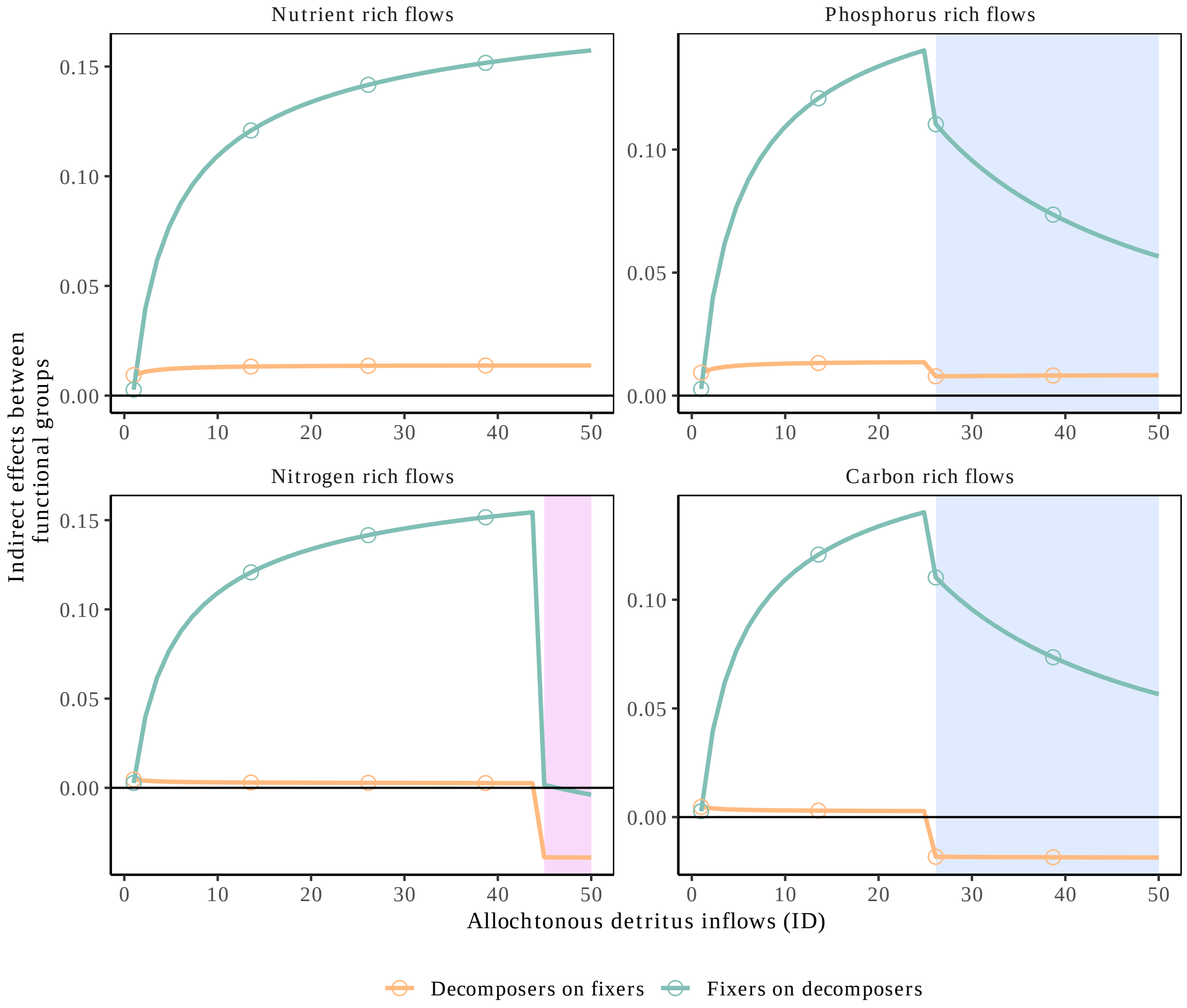


Figure S8: **Allochthonous inflows destabilize indirect effects between functional groups.** Indirect effects between decomposers and fixers along the gradient of allochthonous inflows (ID). The panels correspond to different stoichiometry of the allochthonous flows (low or high P:C and N:C). The blue and purple areas indicate regions where the decomposers are limited by nitrogen or phosphorus respectively (as compared to carbon limitation). Negative values indicate indirect competition while positive values correspond to indirect facilitation.

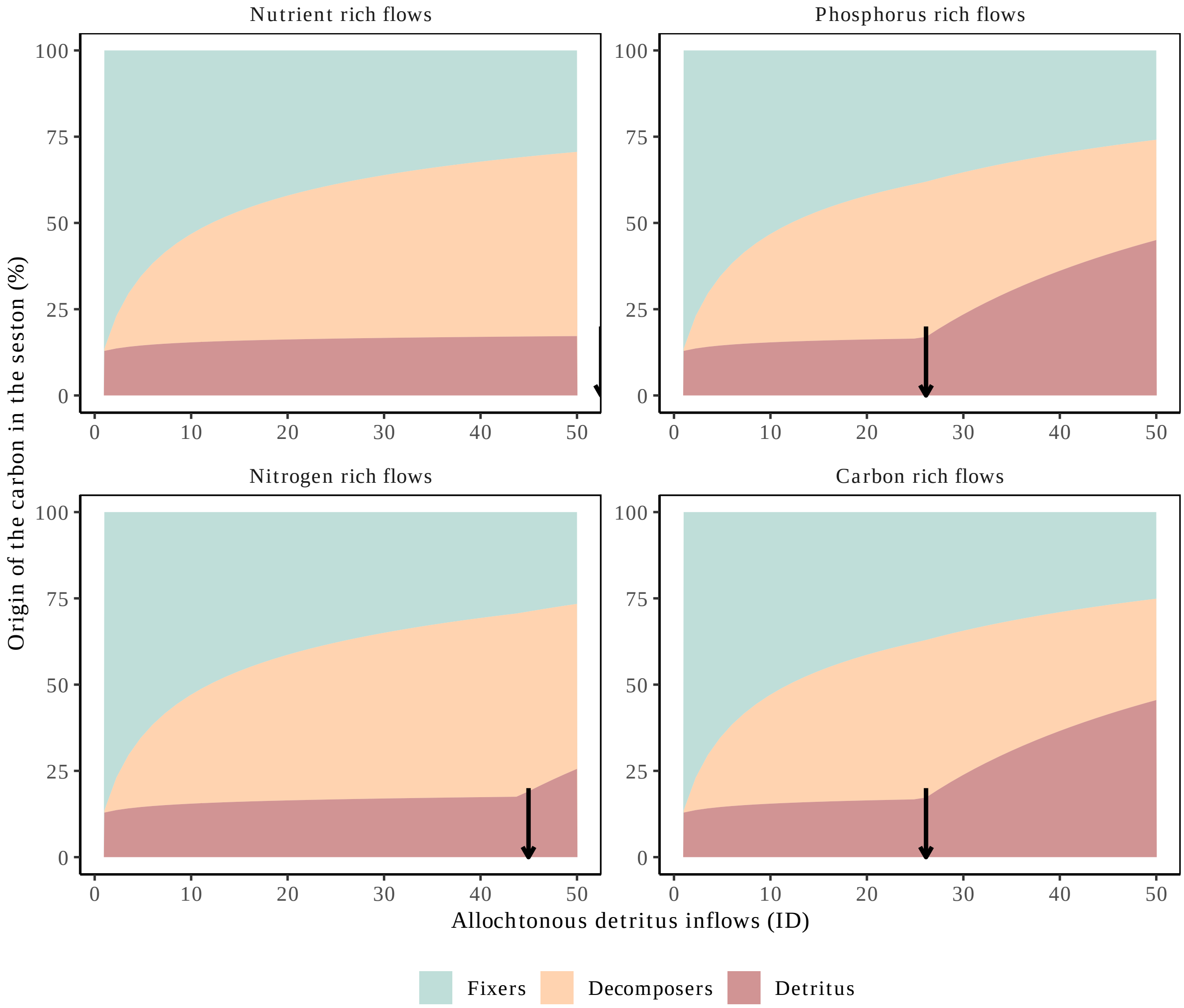


Figure S9: **Decomposer' nutrient limitation hinders the regulation of detritus inflow.** Composition of the carbon in the seston decomposed by the different functional groups of organisms and the detritus. The panels correspond to different stoichiometry of the allochthonous flows (low or high P:C and N:C). The arrow indicate when there is a change in the limitation of decomposers from carbon to nutrients (either nitrogen or phosphorus).

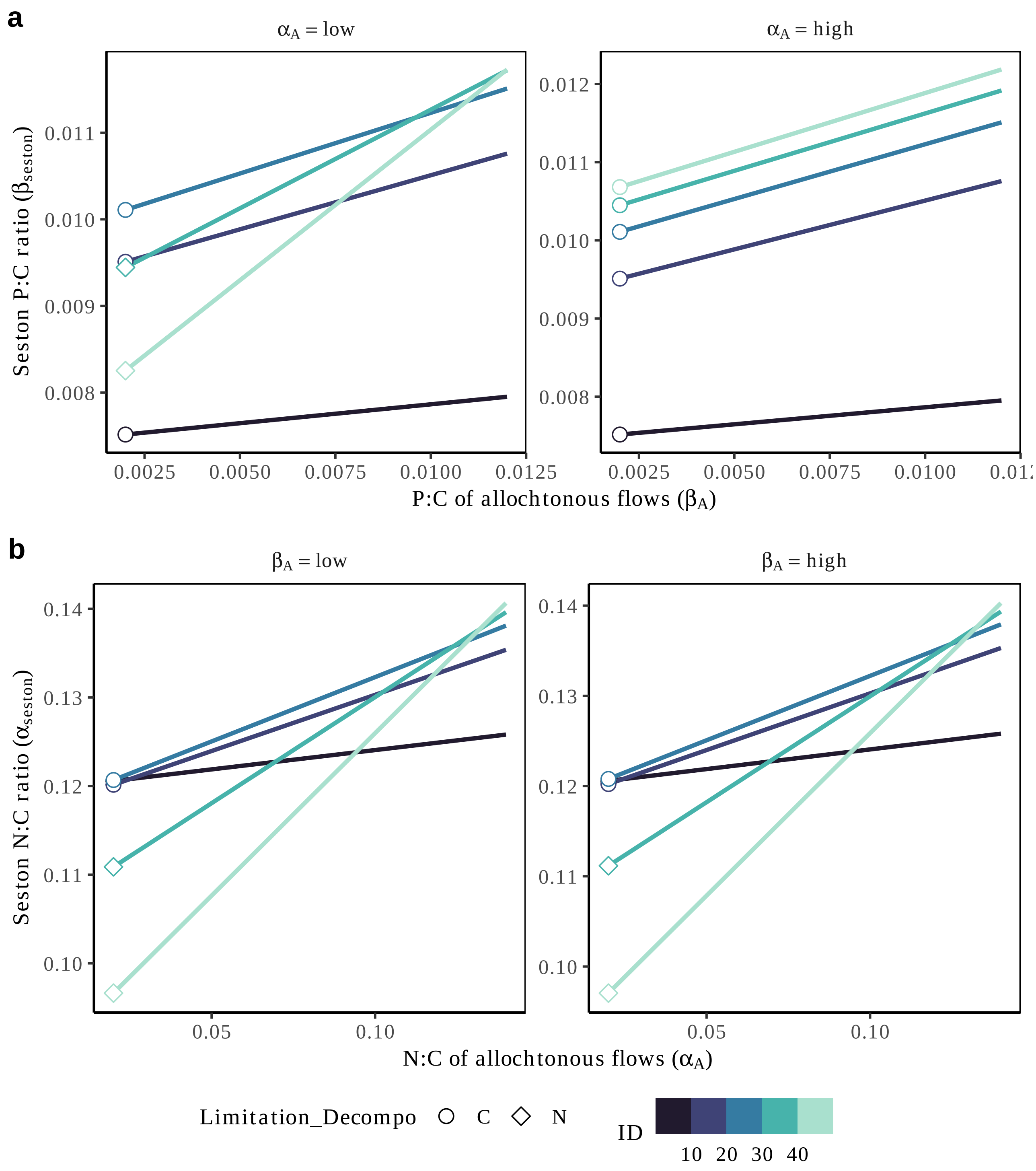


Figure S10: **Sensitivity of the ecosystem to variation in the stoichiometry of allochthonous inflows.**
Ecosystem sensitivity to allochthonous inflow is captured by the plot of the stoichiometry of allochthonous flows (P:C in panel a and N:C in panel B) against the stoichiometry in the seston. The lines are colored by the magnitude of spatial detritus flows and the shape of the points indicate the limitation of decomposers (either N or C limited).

#### *Decomposers and non-fixers*

Setting fixers $F = 0$, we can study the dynamics the two-species model with only decomposers and non-fixers along a gradient of allochthonous inflows quantity and quality.

Increasing allochthonous inflows increases decomposers density. Whether non fixers increase or not in density with allochthonous inflows depends on their stoichiometric quality (Fig. S11). Specifically when $\alpha_B - \alpha_D = \frac{B_N}{B_C} - \frac{D_N}{D_C}$ increases, this means that the N:C of detritus decreases due to poor quality of allochthonous inflows. Because detritus decrease in nitrogen to carbon ratio, decomposers need to take more and more nitrogen, which decreases nitrogen pool as well as non-fixers density (whose dynamics are controlled by nitrogen; Fig. S12). Consequently, the positive indirect effect of decomposers on non-fixers decreases slightly (due to higher indirect competition for nitrogen). Non-fixers by contrast indirectly facilitate decomposers by the production of detritus. The dynamics change when decomposers switch from $C$ to $N$ or $P$ limitation (shaded areas). This change in limitation increase indirect competition between non-fixers and decomposers, limits decomposer growth, and decreases the indirect facilitation between functional groups, and can lead to indirect competition for resources when decomposers are $N$ limited (Fig.XXXb).

As explained in the main text, change in limitation of decomposers was also associated with loss of regulation of detritus inflows (Figs. S13,S14).

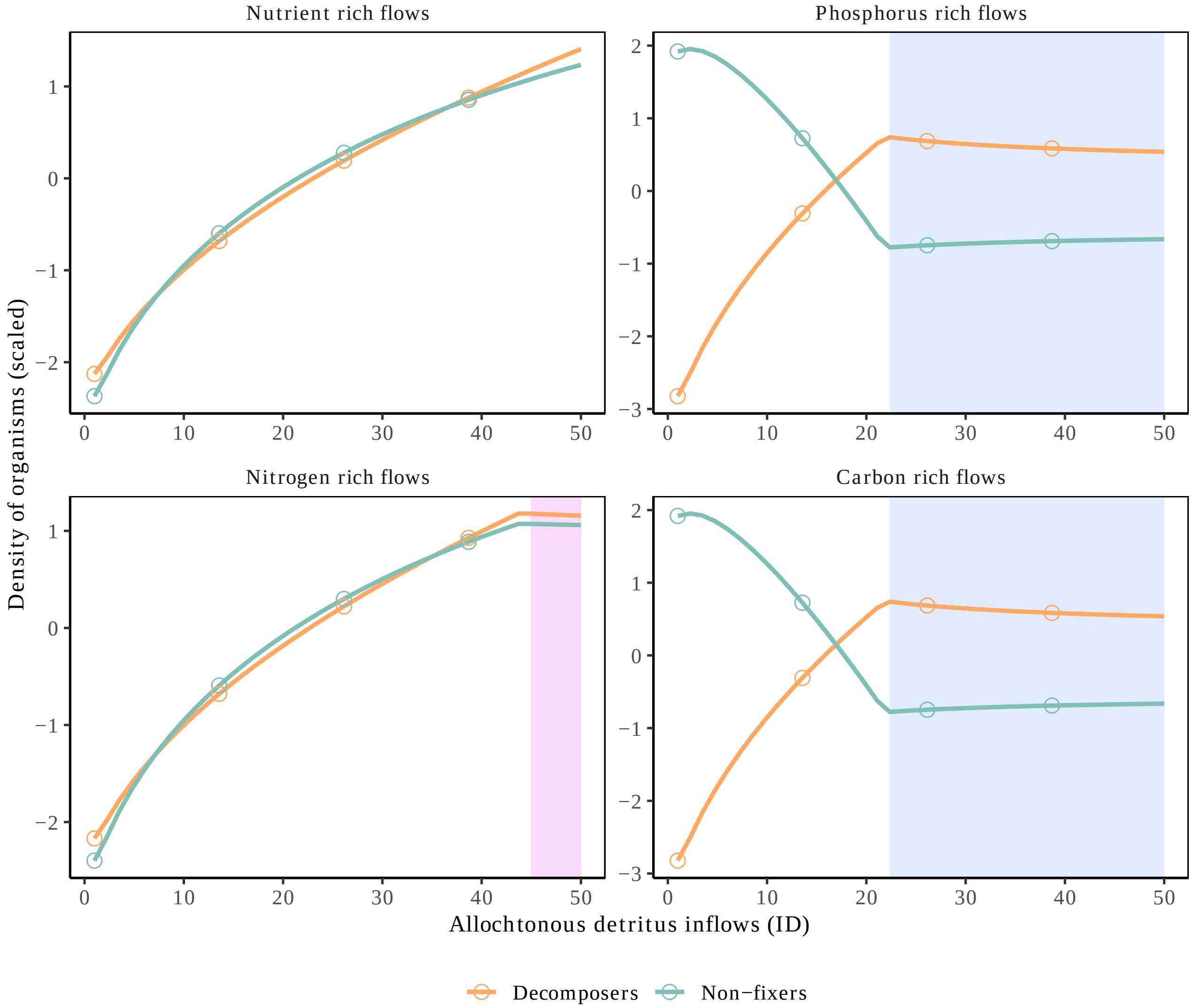


Figure S11: **Dynamics of non-fixers and decomposers along a gradient of allochthonous inflows with varying quality (facets).**
Scaled densities of decomposers and non-fixers (in carbon unit) along the gradient of allochthonous inflows (ID). The panels correspond to different stoichiometry of the allochthonous flows (low or high P:C and N:C). The blue and purple areas indicate regions where the decomposers are limited by nitrogen or phosphorus respectively (as compared to carbon limitation). The densities are scaled to allow comparison of the dynamics along the gradient of ID.

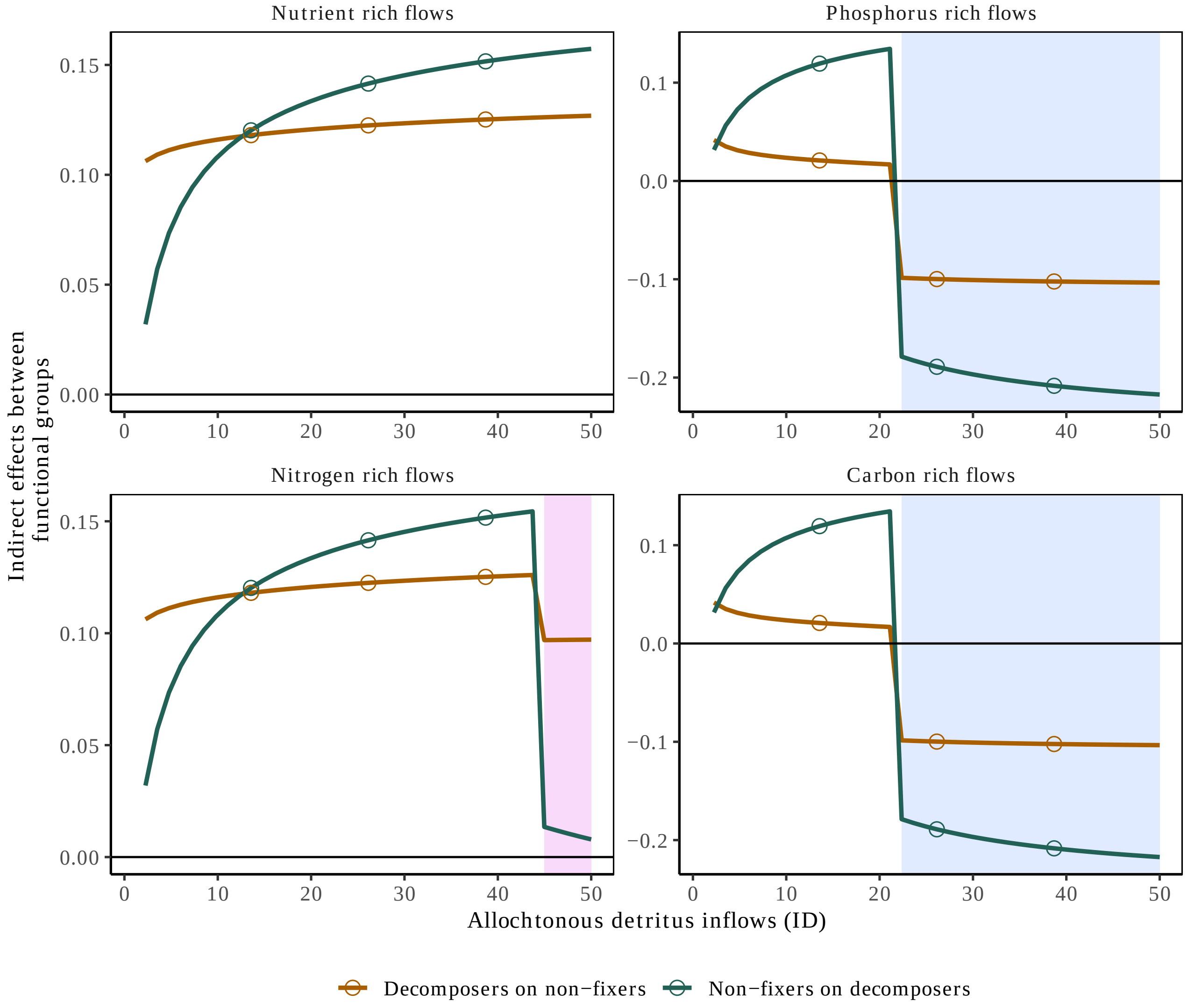


Figure S12: **Allochthonous inflows destabilize indirect effects between functional groups.**
Indirect effects between decomposers and non-fixers along the gradient of allochthonous inflows (ID). The panels correspond to different stoichiometry of the allochthonous flows (low or high P:C and N:C). The blue and purple areas indicate regions where the decomposers are limited by nitrogen or phosphorus respectively (as compared to carbon limitation). Negative values indicate indirect competition while positive values correspond to indirect facilitation.

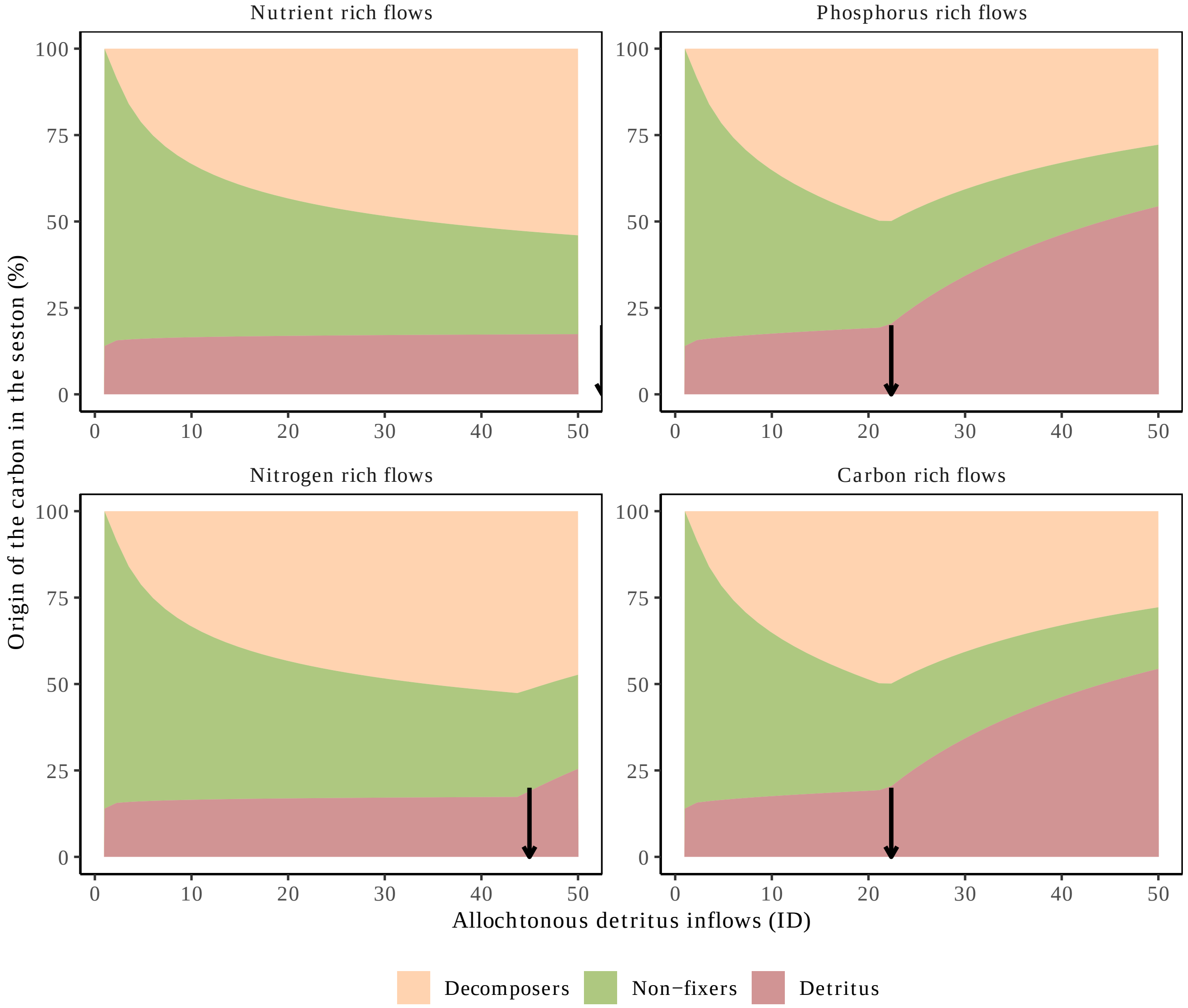


Figure S13: **Decomposer' nutrient limitation hinders the regulation of detritus inflow.** Composition of the carbon in the seston decomposed by the different functional groups of organisms and the detritus. The panels correspond to different stoichiometry of the allochthonous flows (low or high P:C and N:C). The arrow indicate when there is a change in the limitation of decomposers from carbon to nutrients (either nitrogen or phosphorus).

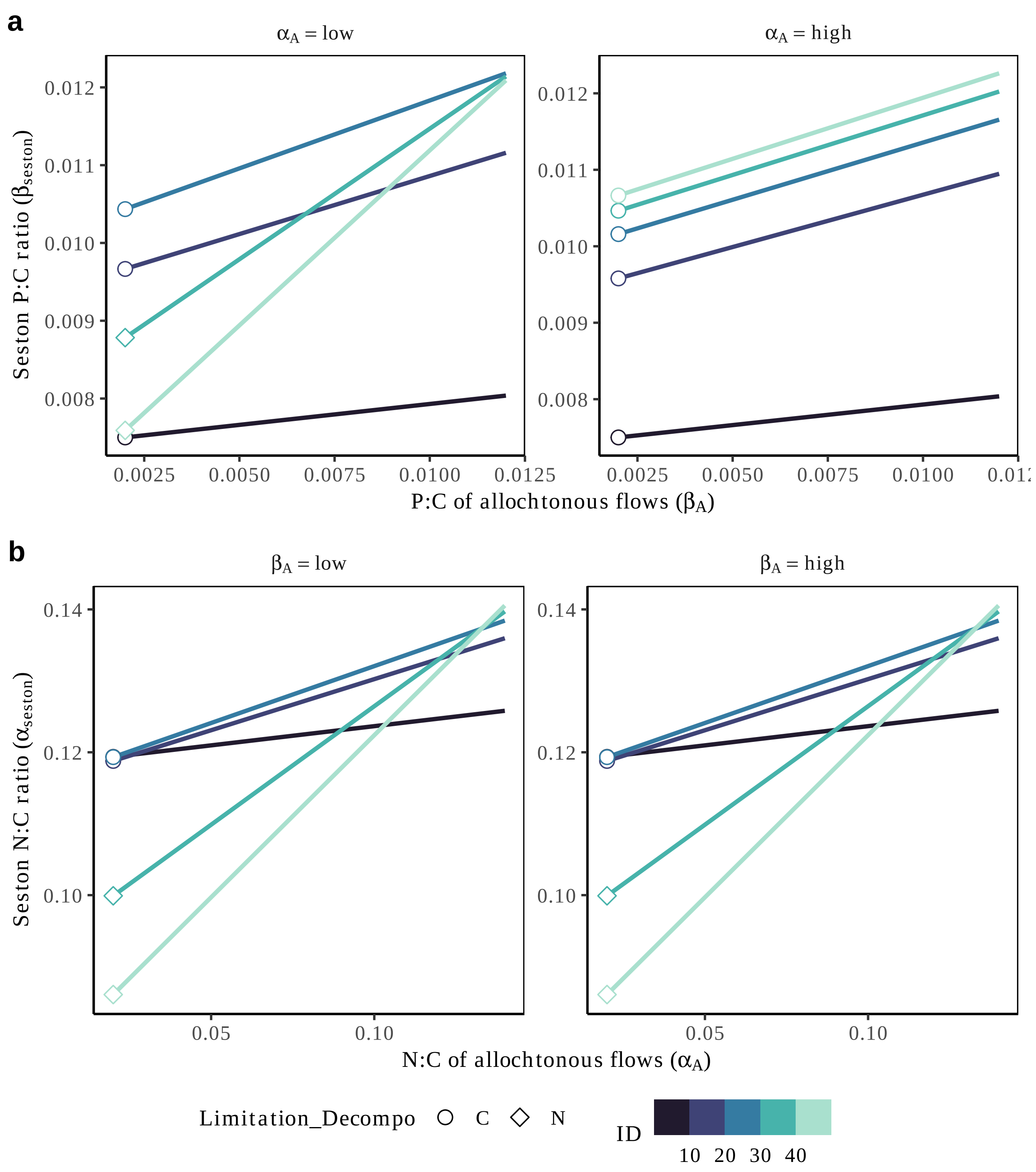


Figure S14: **Sensitivity of the ecosystem to variation in the stoichiometry of allochthonous inflows.**
Ecosystem sensitivity to allochthonous inflow is captured by the plot of the stoichiometry of allochthonous flows (P:C in panel a and N:C in panel B) against the stoichiometry in the seston. The lines are colored by the magnitude of spatial detritus flows and the shape of the points indicate the limitation of decomposers (either N or C limited).

### Appendix S4: Time-varying stoichiometry of allochthonous flows

In this last Appendix, we explore the case of seasonal changes in the stoichiometry of allochthonous inflows. We test two different time-varying changes: a seasonal dynamic of P:C ratio of allochthonous flows and a pulse (therefore transient) of allochthonous flows with high P:C ratio. To model the seasonal dynamics, we only assumes that $\beta_{A,\text{season}} = beta_A(sin(t) + 1)$ so that it can double its value for during certain period of time. This can model the effect of seasonal migration of wildebeest or salmon that enrich aquatic systems with nutrient rich flows (Rüegg et al., 2011; Subalusky et al., 2017). The results are presented in Fig. S15

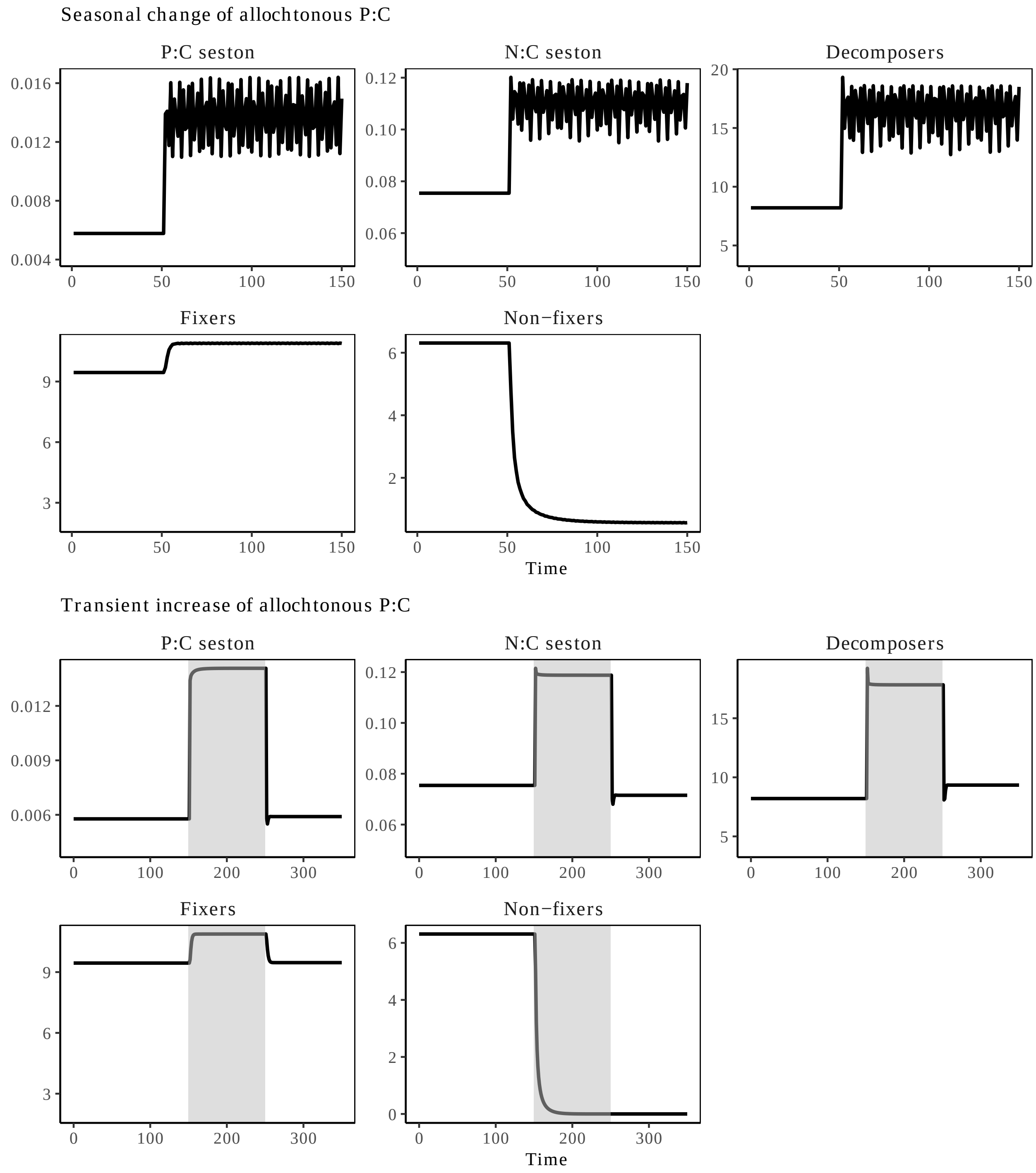


Figure S15: **Example of seasonal change in the C:N:P elemental ratios in the seston.** We show the changes in the elemental ratios and the density of the different functional groups in response to a seasonal dynamic of the P:C ratio of allochthonous flows (a) and a transient increase in the P:C ratio of allochthonous detritus flow (grey area; b). Consequently, we see that the elemental ratios of the seston change a lot through time due to change in the level of heterotrophy, and in the ratio of allochthonous inflows.

### Supplementary material references


Rüegg, J., Tiegs, S. D., Chaloner, D. T., Levi, P. S., Tank, J. L. & Lamberti, G. A. (2011). Salmon subsidies alleviate nutrient limitation of benthic biofilms in southeast Alaska streams. Canadian Journal of Fisheries and Aquatic Sciences, 68, 277–287.

Subalusky, A. L., Dutton, C. L., Rosi, E. J. & Post, D. M. (2017). Annual mass drownings of the Serengeti wildebeest migration influence nutrient cycling and storage in the Mara River. Proceedings of the National Academy of Sciences, 114, 7647–7652.
